# Mechanical regulation of substrate adhesion and de-adhesion drives a cell contractile wave during tissue morphogenesis

**DOI:** 10.1101/2023.05.02.539070

**Authors:** Claudio Collinet, Anaïs Bailles, Thomas Lecuit

## Abstract

During morphogenesis tissue-scale forces drive large-scale deformations, yet how these forces arise from the local interplay between cellular contractility and adhesion is poorly understood. In the posterior endoderm of *Drosophila* embryos, a self-organized tissue-scale wave of actomyosin contractility and cell invagination is coupled with adhesion to the surrounding vitelline membrane to drive the polarized tissue deformation. We report here that this process emerges at the subcellular level from the mechanical coupling between Myosin-II activation and sequential adhesion/de-adhesion to the vitelline membrane. At the wavefront, integrin focal complexes anchor the actin cortex to the vitelline membrane and promote activation of Myosin-II, which in turn enhances adhesion in a positive feedback loop. Subsequently, upon detachment, cortex contraction and advective flow further amplify Myosin-II levels. Prolonged contact with the vitelline membrane increases the duration of the integrin-Myosin-II feedback, integrin adhesion and thus slows down cell detachment and wave propagation of the invagination. Finally, we show that the angle of cell detachment changes as a function of the strength of adhesion and modifies the tensile forces required for detachment to maintain wave propagation. This illustrates how the tissue-scale wave arises from subcellular mechanochemical feedbacks and tissue geometry.

## Introduction

During development active forces due to cell contractility and adhesion drive dramatic tissues shape changes (Lecuit and Lenne, 2007; Lecuit et al., 2011). Spatial and temporal control over the processes generating such forces is essential for robust morphogenesis, whose outcome additionally depends on the mechanical and geometrical environment of the tissue such as its rigidity and the specific shape of its boundaries (Collinet and Lecuit, 2021). It is well established that patterns of gene expression coordinate cell adhesion and contractility in space and time (reviewed in (Collinet and Lecuit, 2021; Gilmour et al., 2017)). For instance, in many organisms the polarized recruitment of non-muscle Myosin-II (MyoII) at the medio-apical surface of epithelial cells, which induces tissue bending and invagination, is controlled by specific transcription factors. Their restricted expression within the tissue regionalizes the activity of specific signaling pathways which culminates in the localized activation of MyoII (Collinet and Lecuit, 2021; Costa et al., 1994; Gilmour et al., 2017; Kolsch et al., 2007; Lee et al., 2006; Manning et al., 2013; Marston et al., 2016; Plageman et al., 2010). Similarly, planar polarized recruitment of MyoII and regulation of cell adhesion molecules at cell-cell contacts depends on specific transcriptional patterns and leads to polarized cell intercalation driving tissue extension (Bertet et al., 2004; Blankenship et al., 2006; Collinet and Lecuit, 2021; Irvine and Wieschaus, 1994; Levayer et al., 2011; Nishimura et al., 2012; Pare et al., 2014; Shindo and Wallingford, 2014; Zallen and Wieschaus, 2004). Besides regulation by transcriptional programs, cell contractility and adhesion display remarkable interplay and self-organization. Many features of contractility such as actomyosin pulses, flows, and waves can only be explained by local stochastic interactions and mechanochemical feedback (Bailles et al., 2019; Bement et al., 2015; Munjal et al., 2015; Nishikawa et al., 2017). These feedback interactions often involve adhesion molecules such as E-cadherin (E-cad) and integrins which are well known to regulate and in turn be regulated by actomyosin contractility (Iskratsch et al., 2014; Kanchanawong and Calderwood, 2023; Leckband and de Rooij, 2014; Lecuit et al., 2011). As such, the local interplay between adhesion and contractility can spatially orient forces. For instance, in *Drosophila* embryos anchorage to E-cad junctions has been shown to bias actomyosin flows and contractions (Levayer and Lecuit, 2013). In turn, the direction of the contractile forces applied to such junctions (producing tensile or shear stress) regulates E-cad levels (Kale et al., 2018). At a larger scale also adhesion to fixed substrates such as the cuticle or the embryonic vitelline membrane has been shown to play an important role in morphogenesis by spatially regulating forces and directing tissue flows (Bailles et al., 2019; Collinet and Lecuit, 2021; Etournay et al., 2015; Munster et al., 2019). Thus, tissue dynamics during morphogenesis emerge from the interplay between contractile and adhesive forces whose spatial-temporal distribution is not only encoded genetically but it is also self-organized and emerges from local mechanochemical feedbacks and the geometry of the system. In this context how the large-scale forces directing tissue flows emerge from the local interplay between cellular contractility and adhesion and how this is regulated by the system’s geometry is poorly understood. Here, we address this during morphogenesis of the posterior endoderm in *Drosophila* embryos, where a wave of actomyosin contractility is coupled to adhesion to the vitelline membrane and propagates in a self-replicating manner (Bailles et al., 2019).

During gastrulation, the posterior endoderm invaginates and a undergoes a polarized flow towards the anterior, thereby driving the initial phase of germ-band extension (Fig.1a top). Here a genetic program induces the initial apical recruitment of MyoII and tissue bending at the posterior of the embryo (Bailles et al., 2019). The recruitment of apical MyoII in this highly-curved region of the embryo leads to a polarized flow of the tissue towards the flatter dorsal side of the embryo, where there is a lower cost to bending (Gehrels et al., 2023). Thus, both genetic and geometric information underlie the onset of polarized flow (Gehrels et al., 2023). After this initial phase where the symmetry is broken and an initial invagination is generated the polarized flow and invagination continues driven by a tissue-level mechanochemical wave of Rho1/MyoII activation propagating from posterior to anterior along the AP-axis from one cell to the next across the embryonic dorsal epithelium (Bailles et al., 2019). A tissue-level characterization (Bailles et al., 2019) emphasized the discrete nature of this process whereby MyoII is activated with rapid kinetics across the entire apical surface of the cells and where activation of MyoII occurs in a given cell at position n only after invagination of the cell at position n-1, suggesting a cell-to-cell relay mechanism. This relay mechanism can be described by a cycle of 3D cell deformations associated with the wave propagation (Fig.1a bottom). The cycle consists of 4 key steps: first, cells anterior to the invaginating furrow are compressed basally and lifted upwards, bringing them in contact with the overlaying vitelline membrane, acting like a rigid substrate that envelops the embryo. Second, the cells spread their apico-lateral surface against the vitelline membrane and adhere via integrins (Bailles et al., 2019; Munster et al., 2019). Third, when cells reach the edge of the advancing invaginating furrow, MyoII contractility is activated in an integrin-dependent manner. Fourth, following MyoII activation cells detach from the vitelline membrane and invaginate into the furrow. This invagination relays the cycle to the next round of cells and subsequent propagation of MyoII activation and cell invagination. Thus, a wave of adhesion, MyoII activation and detachment sustains the persistent movement of the posterior endoderm (Bailles et al., 2019).

**Figure 1:**
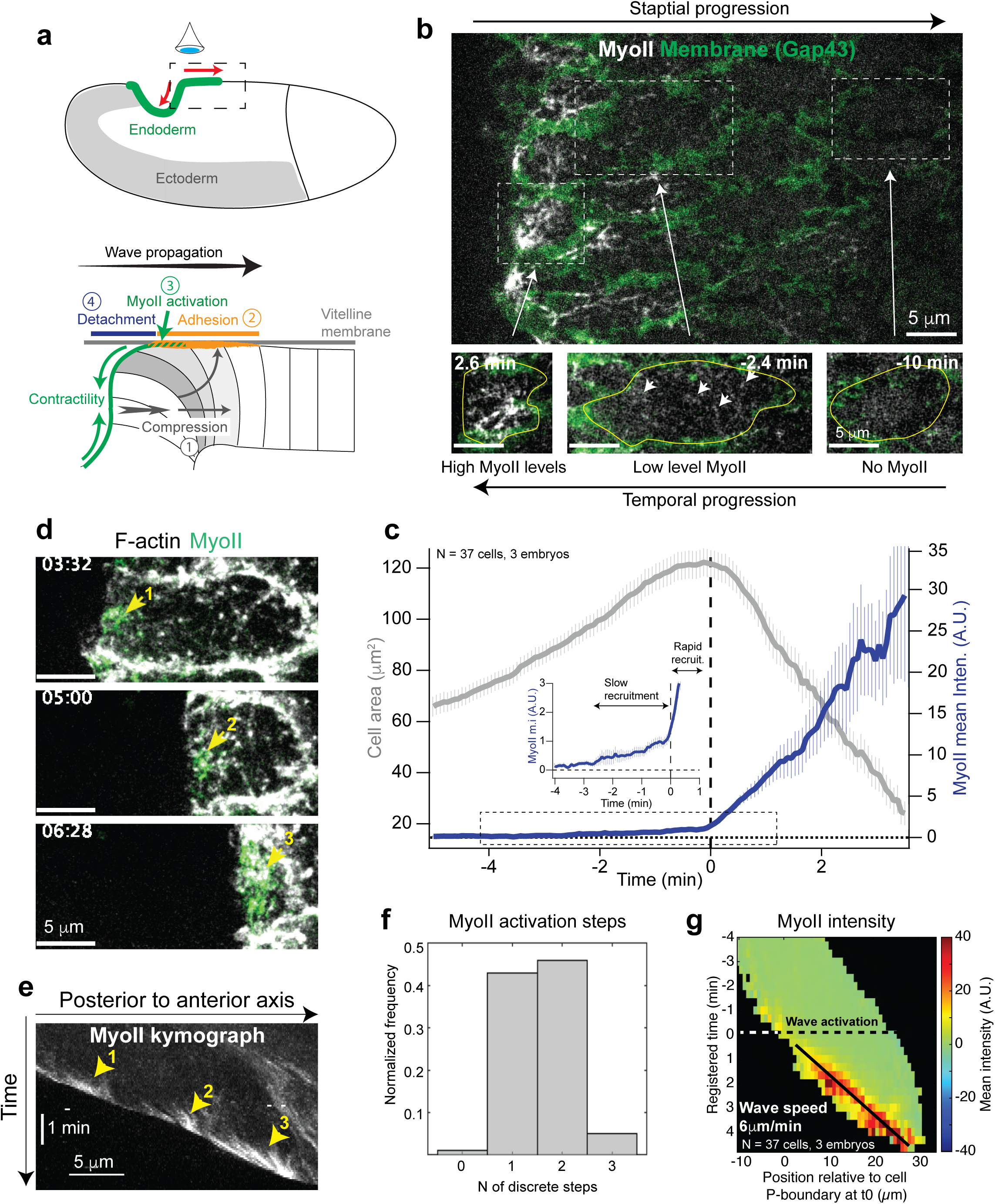
Detailed characterization of MyoII recruitment in cells. (**a**) Cartoon illustration of wave propagation during invagination of the posterior endoderm in *Drosophila* embryos. Top: An embryo where the posterior endoderm is labelled in green and the lateral ectoderm in grey. The boxed region is the region of imaging. The red arrows indicate the directions of wave propagation (to the right) and tissue invagination (to the bottom left). Bottom: Cartoon model of the 3D cycle of cell deformation during wave propagation. In grey: three cells at different stages of the cycle. The numbers indicate the different steps of the cycle. (**b**) High resolution imaging of MyoII recruitment in cells of the propagation region. Top: a larger view on the propagation zone from the invaginating furrow towards the anterior. Bottom: stills from a single cell over time. The boxed regions represent the approximate position of the cell relative to the advancing invaginating furrow. The cell contours are labelled in yellow and the white arrows indicate individual MyoII speckles typical of the slow recruitment phase. (**c**) Quantifications of MyoII mean intensity and cell area over time in cells of the propagation zone. The inset is a zoomed view of the MyoII time trace in the dashed box region. Time 0 is defined for each cell as the onset of rapid MyoII recruitment (see methods). Mean ± s.e.m between cells, N=37 cells from 3 embryos. (**d**) Stills from time lapse of a cell of the propagation zone at the stage of rapid MyoII recruitment. The yellow arrows indicate sequential individual steps of MyoII recruitment just before local cortex detachment. (**e**) Kymograph of MyoII of the cell in **d**. The yellow arrows indicate the sequential steps of MyoII recruitment. (**f**) Histogram of the number of MyoII activation steps in cells of the propagation zone. N=95 cells, 5 embryos. (**g**) Kymograph heatmap of MyoII mean intensity in cells during wave propagation. N=37 cells, 3 embryos. Cells were registered in time as in **c** and in space by positioning their posterior boundary (P-boundary) at x=0 at time t=0 (see also Suppl. Fig. 1c-d).

Although this model explains how MyoII contractility is linked to the anterior movement of the tissue, several key questions remain unanswered. For instance, it is unclear by which mechanism MyoII activation occurs in contact with the vitelline membrane and how integrins regulate this. Furthermore, it is unclear how MyoII contractility in contact with the vitelline membrane and within the furrow interacts with adhesion mediated by integrins to detach the cells and relay the cycle. To address these questions here we studied the detailed dynamics of MyoII and integrins recruitment in contact with the vitelline membrane and how the cycle of cell attachment/detachment to it mediates wave propagation.

## Results

### A subcellular wave of MyoII propagates within cells

To better characterize the process of MyoII recruitment we imaged cells of the *Drosophila* posterior endoderm during the process of wave propagation at high spatial and temporal resolution. We expressed the Actin binding domain of Utrophin (Utr-ABD) fused to EGFP or Gap-43 fused to mCherry, to label F-actin or cell membranes respectively, and the MyoII Regulatory Light Chain (MRLC) fused to mCherry or EGFP, to label MyoII. We observed two phases of MyoII recruitment in cells during wave propagation. First, at a 1-2 cell distance from the furrow when cells adhered to the vitelline membrane, we observed MyoII recruitment at low levels (Fig.1b-c, Suppl. Fig.1a and Movie 1). This phase was characterized by the appearance of transient MyoII speckles, reminiscent of individual minifilaments, across the entire apical surface of the cell (Fig.1b, Movie 1 and Suppl. Fig.1b). Then, when cells were near the advancing border of the invaginating furrow, MyoII recruitment significantly accelerated (Fig.1b-c and Suppl. Fig.1a) and occurred in discrete steps; bright clusters of MyoII appeared at the posterior of the cell just before local detachment from the vitelline membrane (Fig.1d-e, Suppl. Fig.1b and Movie 2). Several clusters appeared per cell as the cell detached (Fig.1f), first near the cell posterior then more anteriorly, indicating that high MyoII recruitment propagates inside the cell (Fig.1d-e and Movie 2).

We tracked and registered cells over time and space (see methods and Suppl. Fig.1c-e) and plotted the local mean intensity of MyoII as a function of time and distance from the posterior cell boundary at time 0 (Fig.1g and Suppl. Fig.1d). Using this representation, we visualized the intracellular propagation of MyoII in the fixed reference frame of the embryo and estimated its velocity to ∼6 µm/min, which is similar to the observed velocity of the tissue level-wave in the same frame of reference (Bailles et al., 2019).

Thus, a subcellular propagation of MyoII concentration and cell detachment propagate within cells as a wave with similar kinetics to those of the tissue-level wave, suggesting that similar mechanisms underlie the two and that the latter might be a larger-scale manifestation of the former. This led us to further investigate the mechanism driving the subcellular wave propagation.

### Integrin adhesion to the vitelline membrane immobilizes the apical actin cortex prior to wave propagation

Previous work suggested that cells engaged in apical frictional coupling and adhesion to the vitelline membrane before activating MyoII (Bailles et al., 2019; Munster et al., 2019). We thus tested whether the medio-apical actin cortex adhered to the overlying immobile vitelline before propagation of the intracellular wave of MyoII. The Utr(ABD)::EGFP probe labelled F-actin filaments and concentrated actin forming bright spots in the apico-medial cortex (Dehapiot et al., 2020) (Fig. 2a-b and Movie 3) and we tracked their movement in cells during wave propagation using the KLT feature tracker algorithm (Dehapiot et al., 2020; Lucas and Kanade, 1981; Tomasi and Detection, 1991). We found that before cell spreading onto the vitelline membrane, actin filaments and bright particles in the medio-apical cortex moved anteriorly together with cell boundaries (Fig.2a, Movie 3 and Fig.2c upper part of the kymograph) and displayed an average positive velocity of 1.65±0.02 s.e.m. µm/min (Fig.2e). Subsequently, upon spreading onto the vitelline membrane and before cell detachment from the posterior of each cell, actin particles significantly slowed down (Fig.2b, Movie 3 and Fig.2c bottom part of the kymograph) and displayed an average velocity of 0.20±0.02 s.e.m. µm/min (Fig.2e), indicating an immobilization of the apical actin cortex. Notably the MyoII wave was triggered after contact with the vitelline membrane and propagated across the immobilized actin cortex (Fig.2c).

**Figure 2:**
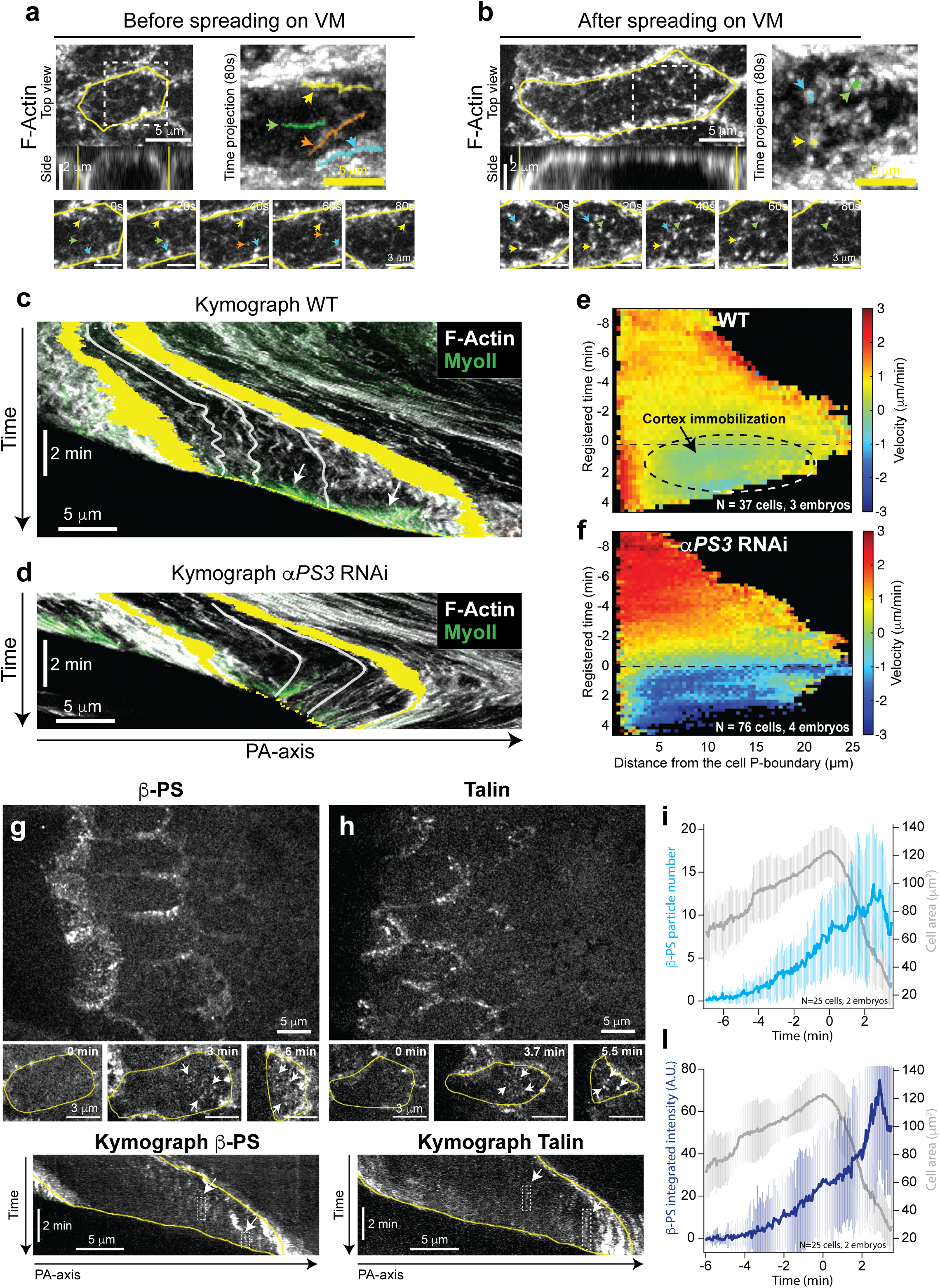
Actin cortex immobilization and Integrin recruitment during wave propagation. (**a-b**) Apical actin cortex during wave propagation. On the left: Top and side views before (**a**) and after (**b**) spreading onto the vitelline membrane. The cell contour is in yellow. At the bottom: snapshots of the boxed regions in the top images with an interval of 20s. The colored arrowheads indicate individual actin speckles followed over time. On the right: a time projection over 80s of the boxed regions in the top images. The tracked actin speckles are indicated by the colored arrowheads and their trajectories by the semitransparent colored lines. (**c-d**) Kymographs of F-actin (in grey) and MyoII (in green) along the PA-axis in a WT (**c**) and a *αPS3* RNAi injected embryo (**d**). The anterior and posterior boundaries of a cell are highlighted in yellow. The white semitransparent lines indicate traces of actin speckles. (**e-f**) Heatmap kymographs of actin particles velocities for WT (**e**) and *αPS3* RNAi injected embryos (**f**). Particle velocities were measured in the referential of the embryo and the data were plotted relative to the moving P-boundary of the cell. The dashed oval region indicates cortex immobilization estimated as the region where velocities are close to 0. N=37 cells from 3 embryos for WT and, 76 cells from 4 embryos for *αPS3* RNAi. (**g-h**) Stills of β-PS (**g**) and Talin (**h**) in cells of the posterior endoderm during wave propagation. Top: larger scale view. Middle: close-ups on a single cell over time. Yellow: cell contours. White arrows: medio-apical puncta forming focal contacts. Bottom: Representative kymographs along the PA-axis of individual cells. White arrows indicate immobile puncta forming focal contacts (boxed by the white dashed line) in the medio-apical region of the cell. The A-and P-boundaries of the cell are in yellow. (**i-l**) Time traces of cell area together with the number of β-PS particles per cell (**i**) or their total integrated intensity (**l**). Mean ± SDs between cells. N=25 cells from 2 embryos. In **e**, **f**, **i** and **l** time 0 is the time of wave activation (rapid MyoII recruitment, see methods).

The immobilization of the actin cortex prior to MyoII recruitment is reminiscent of the formation and maturation of nascent adhesions by integrins during cell migration (Kanchanawong and Calderwood, 2023). To test whether the apical cortex immobilization was due to integrin-mediated cell adhesion to the vitelline membrane we depleted by RNAi the αPS3 integrin, known to be involved in the adhesion of the posterior endoderm to the vitelline membrane (Bailles et al., 2019; Munster et al., 2019). This changed the pattern of cell movement (Fig.2d) and, consequently, of the actin particle velocities (Fig.2f). Consistent with our previous findings (Bailles et al., 2019), in *αPS3* RNAi embryos, cells first underwent a phase of fast movement towards the anterior, failed to immobilize against the vitelline membrane and eventually moved backwards towards the invaginating furrow (Fig.2d). The actin particles moved together with the cell and displayed strong negative velocities (i.e. directed towards the posterior) indicating that in the absence of αPS3 the cortex is not immobilized. We conclude that the actin cortex immobilization prior to subcellular MyoII wave propagation depends on integrin-mediated adhesion to the vitelline membrane.

These findings led us to analyze the dynamics of integrins and of integrin-associated proteins (IAPs) to better understand integrin mediated adhesion during this process. In *Drosophila* the βPS integrin is the main β-Integrin (out of only 2 present in the genome) and is the only one present in the embryo at gastrulation stages (Yee and Hynes, 1993). βPS forms complexes with all of the 5 α-subunits present in the *Drosophila* genome (Brown et al., 2000). In the embryo βPS and Talin, an essential IAP present nearly at all stages of integrin adhesions formation and maturation (Iskratsch et al., 2014; Kanchanawong and Calderwood, 2023; Klapholz and Brown, 2017; Nayal et al., 2004), are ubiquitous and deposited maternally (Brown et al., 2002; Sawala et al., 2015b) while only 3 of the 5 α-subunits are expressed in restricted dorsal and ventral regions at gastrulation stages(Sawala et al., 2015b) (Suppl. Fig.2a). To visualize integrins we used knock-in lines of the βPS-integrin and of Talin fused to EGFP (Klapholz et al., 2015) and Y-Pet (Lemke et al., 2019) respectively. Using these tools, we found that βPS accumulated in bright intracellular vesicles and at the plasma membrane in bright puncta, reminiscent of focal complexes (Kanchanawong and Calderwood, 2023), only in a dorsal posterior region, where αPS3 is expressed. In other regions of the embryos, βPS remained in faint intracellular structures resembling the endoplasmic reticulum (ER) (Suppl. Fig.2b), suggesting that delivery to the cell surface is blocked as expected from a missing α-chain. Similarly, Talin showed a cortical accumulation in cells in the same dorsal posterior region while elsewhere it was found in the cytoplasm (Suppl. Fig.2c). This was consistent with the localization of βPS and suggests that during gastrulation βPS forms functional integrin heterodimers involved in substrate adhesion only where αPS3 is expressed. Consistent with this, in *αPS3* RNAi embryos βPS lost its plasma membrane localization in puncta (Suppl. Fig.2d). We conclude that αPS3 is key to induce the formation of focal complexes in the posterior endoderm where the wave of MyoII activation and cell invagination propagates.

To quantitatively analyze focal complex dynamics during wave propagation we imaged the apical planes of cells in contact with the vitelline membrane at high spatial and temporal resolutions. We found that during cell apical spreading onto the vitelline membrane the newly formed βPS and Talin puncta in the medio-apical side of the cell were immobile in the referential of the vitelline membrane (Fig.2g-h and Movies 4-5). The number of βPS puncta increased more than 10-fold over time before cell detachment occurred (Fig.2i), while the average size of puncta and average intensity increased less (2-3 fold, Suppl. Fig.3). Thus, the global increase in total integrated βPS intensity mostly stemmed from an increase in the number of puncta (Fig.2l and Suppl. Fig.3).

**Figure 3:**
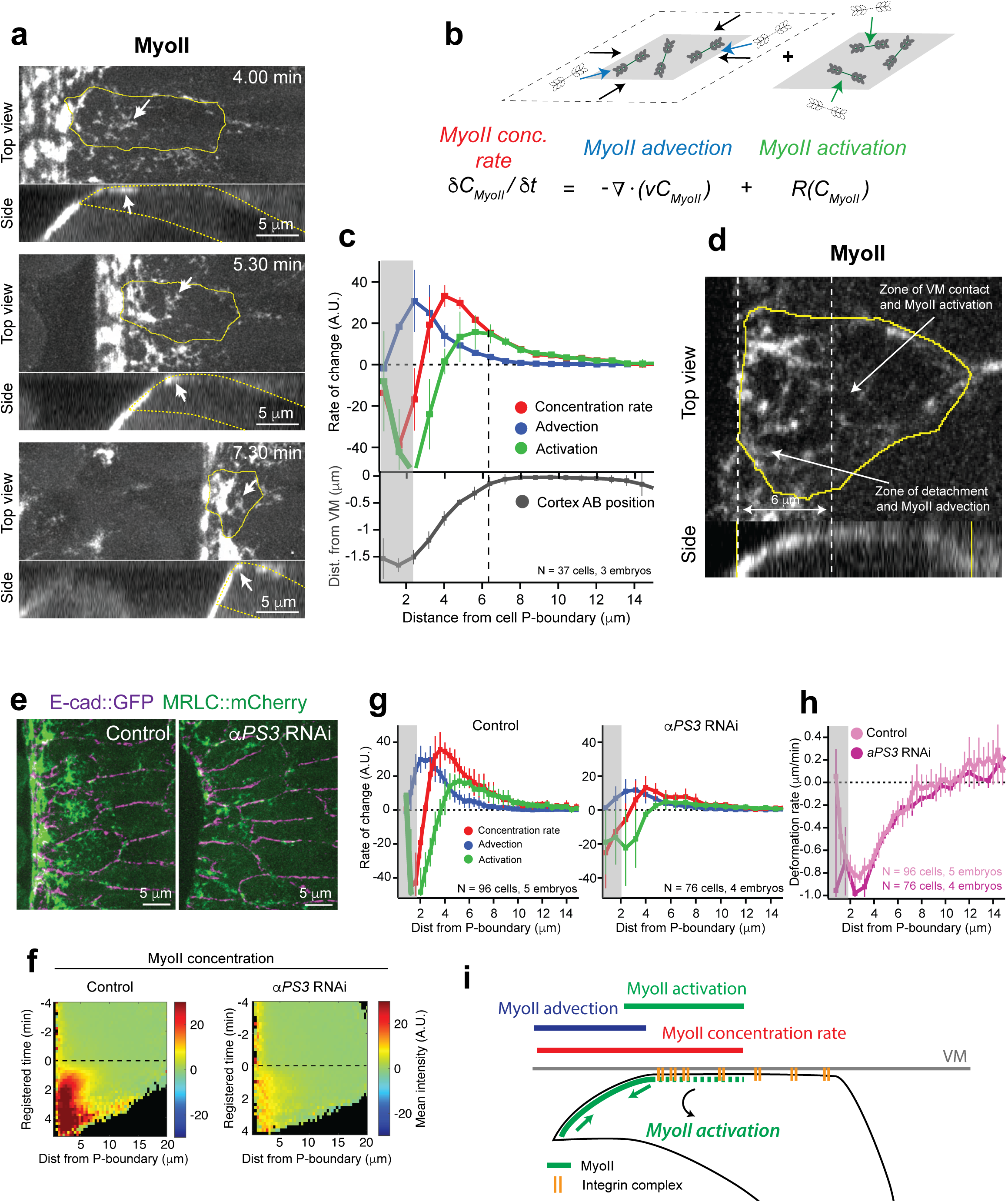
Integrin-dependent MyoII activation in contact with the vitelline membrane. (**a**) Representative stills (top and side views) of MyoII propagation in a cell. The white arrows indicate foci of MyoII initial recruitment in contact with the vitelline membrane. In yellow are the cell contours. (**b**) Equation of local mass balance for MyoII in an coarse-grained moving piece of cortex. The contribution of MyoII activation (R_MyoII_) and advection are stated explicitly. The cartoons on top of the equation terms represent a change in MyoII local concentration by advection with cortex contraction (left cartoon) and by the turnover rate of MyoII minifilaments binding to the cortex (right cartoon). (**c**) Average spatial profiles of MyoII rates along with the cortex apico-basal position in WT cells during wave propagation. The dashed line indicates cortex detachment. Mean ± SDs between embryos. N=3 embryos (37 cells). (**d**) Representative top and side view of MyoII during wave propagation. The zones in contact with and detached from the vitelline membrane are indicated. (**e**) Representative stills of MyoII and E-cad during wave propagation in a control (water injection) and a *αPS3* RNAi embryo. (**f**) Heatmap kymographs of MyoII mean intensity in the indicated conditions. Data are plotted relative to the moving P-boundary of the cells. N=96 cells from 5 embryos for control (water injection) and N=76 cells from 4 embryos for *αPS3* RNAi. (**g-h**) Average spatial profile of MyoII rates (**g**) and actin cortex deformation rate (**h**) in the indicated conditions during wave propagation. In **h** a negative rate indicates local contraction during wave propagation. Mean ± SDs between embryos. N=5 embryos (96 cells) for control (water injection) and N=4 embryos (76 cells) for *αPS3* RNAi. (**i**) Cartoon summary of the spatial distribution of the 3 measured MyoII rates in a cell during wave propagation. VM is the vitelline membrane.

We conclude that apical cell adhesion to the vitelline membrane is organized in focal complexes, whose number increases over time. This immobilizes the underlying actin cortex before activation of the intracellular wave of MyoII.

### MyoII activation occurs in contact with the vitelline membrane and depends on integrins

To better understand how MyoII propagates as a wave inside cells, we analyzed how and when MyoII is recruited with respect to cortex adhesion to and detachment from the vitelline membrane. Qualitative observations from top and side views (Fig.3a and Movie 6) first showed that during wave propagation MyoII clusters appeared in contact with the vitelline membrane and became brighter upon cell detachment (Fig.3a and Suppl. Fig.4b). This suggested that MyoII recruitment might be induced in contact and further increase upon detachment from the vitelline membrane. A local change in MyoII concentration depends on two processes: 1) the transport by advective flows due to cortex contraction of MyoII mini-filaments bound to the actin cortex and 2) the net association/dissociation of MyoII mini-filaments to the actin cortex from the cytoplasmic pool (here referred to as MyoII activation rate, R_MyoII_, which captures turnover and Rho pathway-mediated recruitment). To disentangle the contribution of these two effects during wave propagation, we imaged F-actin together with MyoII. We locally measured the MyoII intensity and used KLT to determine the velocity of the actin cortex (see methods). These measurements allow us to determine the local rates of MyoII concentration change, MyoII advection and R_MyoII_ using mass conservation (Nishikawa et al., 2017; Vallotton et al., 2004) (Fig.3b).

We mapped these quantities within cells during wave propagation and correlated their local distribution with the contact to vitelline membrane (Fig.3c). We found that MyoII advection and activation spatially segregate and contribute to MyoII concentration in different regions of the cell (Fig.3c). In particular, in more anterior regions where the actin cortex is in contact with vitelline membrane (Fig.3d) R_MyoII_ dominated and contributed almost entirely to the rate of MyoII concentration change. More posteriorly, where the cortex was detached from the vitelline membrane (Fig.3d), MyoII advection increased significantly and contributed to further increase the MyoII concentration rate (Fig. 3c) and MyoII levels (Suppl. Fig.4a-b). This suggests that the MyoII wave propagates within the cell with a MyoII activation front that recruits new MyoII mini-filaments in contact with the vitelline membrane. This is followed by further MyoII concentration by advection once the cortex detaches from the vitelline membrane (Fig.3i). The time evolution of the MyoII rates in a fixed region of the cell cortex confirmed this. R_MyoII_ increased first when the cortex was still in contact with the vitelline membrane. Later on, upon cortex detachment from the vitelline membrane, MyoII advection increased and contributed to further increase the rate of MyoII concentration (Suppl. Fig.4e).

Since focal complexes are known to respond to force (Iskratsch et al., 2014; Kanchanawong and Calderwood, 2023; Sun et al., 2016) and regulate Rho signaling and MyoII contractility in other systems (Guilluy et al., 2011; Schiller and Fassler, 2013; Sun et al., 2016), we next investigated how integrins regulate MyoII recruitment and the rates of advection/activation. As expected, depleting αPS3 by RNAi reduced the levels of MyoII during wave propagation (Bailles et al., 2019) (Fig.3e-f and Movie 7). Consistent with this, the rates of MyoII concentration, MyoII advection and R_MyoII_ were also significantly reduced compared to control embryos (Fig.3g). Interestingly however, the rate of cortex deformation (the strain rate) was not significantly different from that of control embryos (Fig.3h), indicating that the reduced MyoII advection and concentration rate stem from a lower initial recruitment of MyoII mini-filaments (which reduces C_MyoII_ and thus MyoII advection, Fig.3b). We thus conclude that the primary effect of depleting integrins is to reduce R_MyoII_ in contact with the vitelline membrane.

These results indicated that during wave propagation MyoII accumulation occurs by two distinct and spatially segregated mechanisms. First, MyoII is activated by integrins in contact with the vitelline membrane where the cortex is immobile and then it further accumulates by advection upon cortex detachment when the cortex contraction is no longer resisted by integrin adhesion (Fig.3I). Notably, upon cortex detachment MyoII promotes its own accumulation in a positive feedback loop (Munjal et al., 2015).

### MyoII contractility controls focal complexes maturation and stability

Since integrins are known to respond to forces exerted by actomyosin contractility (Gauthier and Roca-Cusachs, 2018; Iskratsch et al., 2014; Schiller and Fassler, 2013), we next investigated whether this was the case also for the focal complexes we observed during wave propagation. To that end we imaged MyoII together with the βPS integrin. We observed that βPS-positive puncta in the medio-apical region of the cell formed much earlier than the bright MyoII clusters of the wave (Fig.4a, left and kymographs and Movie 8). Interestingly, shortly before cortex detachment, MyoII clusters formed in proximity of βPS puncta and appeared connected to them, suggesting the MyoII clusters might be anchored to and pulling on the focal complexes (Fig4a, inset and Movie 8).

To further test this possibility, we imaged Vinculin, a protein known to be recruited to focal complexes upon force during their maturation to Focal Adhesions (FAs) (Atherton et al., 2016; Hirata et al., 2014; Kanchanawong and Calderwood, 2023; Sun et al., 2016), together with either βPS or MyoII. As expected, Vinculin labelled E-cadherin junctions (Kale et al., 2018) but it also appeared in bright medio apical puncta in contact with the vitelline membrane (Fig.4b and Movie 9). However, in contrast to βPS, Vinculin puncta appeared only shortly before cell detachment when the bright MyoII clusters of the intracellular wave appeared (Fig.4b, Movie 9 and Suppl. Fig.5a). We thus hypothesized that MyoII clusters of the wave pull on pre-formed focal complexes inducing Vinculin recruitment on these structures. Consistent with this idea, we observed that Vinculin appeared medio-apically later (Fig.4c) at preformed βPS puncta just before cell detachment (Suppl. Fig.5a). Furthermore, a positive cross-correlation of 0.34±0.04 s.e.m between the rates of change of the total fluorescence intensity of Vinculin and MyoII with a time delay of ∼12s indicated that a strong increase in Vinculin signal shortly followed a rapid increase in MyoII (Fig.4d-e).

These data suggest that during wave propagation MyoII contractility is required for focal complex maturation and thus stabilization. To test this directly, we blocked MyoII contractility by injecting a Rok inhibitor (H-1152). This led to the rapid reduction in MyoII concentration at the cortex and arrest in wave progression. It also affected recruitment of βPS at focal contacts: βPS puncta faded away both in the cells at the edge of the furrow where the intracellular MyoII wave propagates (Fig.4f-i and Movie 10) and in the more anterior cells where focal contacts form in contact with the vitelline membrane (Suppl. Fig.5b-d). We conclude that MyoII activity is required for the stabilization and maturation of focal complexes. Wave propagation is thus characterized by positive feedback between integrins and MyoII: integrins are required to activate MyoII and MyoII sustains stabilization of integrin focal contacts. This positive feedback loop operates in contact with the vitelline membrane where the orientation of the actin filaments parallel to the surface induces shear stress on integrin complexes.

### Cortex detachment allows MyoII wavefront propagation and controls the duration of the Integrin-MyoII feedback

Since integrin-MyoII positive feedback operates in contact with the vitelline membrane, we next investigated whether the kinetics of detachment might regulate the speed of the wave and the levels of MyoII and adhesion during wave propagation. We hypothesized that cortex detachment is promoted by tensile forces due to MyoII contractility in the invaginated furrow which are transmitted to the anchoring point at the front of the invagination (Fig.5a left bottom diagram). We thus sought to mechanically interfere with cell detachment in two ways. First, we mildly depleted α-Catenin (α-Cat) in embryos by RNAi. This is known to weaken the connections of the actomyosin cortex to E-Cadherin junctions (Cavey et al., 2008) and thus force integration across the tissue(Martin et al., 2010) and their transmission to the anchoring point at the front of the furrow. Second, using laser-mediated tissue cauterizations (Collinet et al., 2015; Rauzi et al., 2015) we generated immobile mechanical fences 10-12 cells anterior to the moving furrow to introduce a mechanical resistance to the furrow movement and to cell invagination (Bailles et al., 2019) (Movie 11). In both cases we observed a slower anterior progression of the invaginating front compared to control embryos (Fig.5a-b and Movie 12) associated with a slower detachment of the local cell cortex following local MyoII recruitment (Fig.5b-c and Suppl. Fig. 6a, grey curves). Interestingly, we observed that within single cells across different conditions the speed of cortex detachment highly correlated with the speed of the MyoII wave as measured in kymographs (Fig.5b and 5d). This indicates that local detachment is required to propagate the MyoII wavefront within the cell, thus setting the pace of the wave.

The increased time of contact with the vitelline membrane due to the slower detachment is expected to prolong the activity of the integrin-MyoII positive feedback and thereby lead to prolonged MyoII activation, higher MyoII levels and increased accumulation of focal complexes mediating adhesion at the wavefront (Fig. 6a). To test this prediction, we first measured the rate of MyoII concentration, MyoII advection and R_MyoII_ over time in given regions of the cell cortex during wave propagation (as shown in Suppl. Fig.4c-e). In both *α-cat* RNAi and embryos with anterior fences, R_MyoII_ was similar to controls but remained positive for a longer time (Fig.6b and Suppl. Fig.6b, green curves). This was consistent with the observed delayed detachment of the cortex (Fig.6b and Suppl. Fig.6b, grey curves) and explains the higher levels of MyoII in these conditions (Suppl. Fig.6a). Next, we tested if this phenomenon was integrin-dependent. Simultaneous depletion of both α-Cat and αPS3 by RNAi significantly reduced MyoII levels and R_MyoII_ to levels similar to that in the case where only αPS3 was depleted (Fig.6c-e and Movie13). To test if the prolonged integrin-MyoII positive feedback also increased integrin clustering in focal complexes, we imaged βPS together with MyoII in either *α-cat* RNAi or embryos with anterior immobile fences. The local density of βPS and of the βPS-positive puncta increased (Fig.6f-g and Movie 14-15). Interestingly the average size and the average total intensity of the puncta also increased (Suppl. Fig.6c-d), indicating the focal complexes can mature into larger structures such as FAs upon longer time of contact. This led to increased cell adhesion to the vitelline membrane as individual cells frequently resisted detachment inducing an irregular advancing furrow (Fig.6f and Movie 14-15).

We conclude that local cortex detachment allows MyoII wavefront propagation and controls the duration of the integrin-MyoII feedback activation mechanism.

### The angle of detachment changes according to the adhesion strength and modifies de-adhesion tensile forces

Cortex detachment plays a key role in wave propagation as it regulates the speed of the MyoII wavefront and the duration of the integrin-MyoII feedback. We thus investigated how detachment is controlled from the standpoint of the local forces causing integrin de-adhesion from the vitelline membrane (Fig.7a) and considered the role of tissue geometry. At the apical surface in contact with the vitelline membrane MyoII contractility causes shear forces on integrins that reinforce focal complex formation and maturation (Plotnikov et al., 2012; Raz–Ben Aroush and Wagner, 2006) (Fig.4). On the contrary, MyoII contractility in the furrow and the detached posterior of the cell, which is at an angle θ from the vitelline membrane (Fig.7a), exert tensile forces that might cause de-adhesion of integrins and, consequently cell detachment. We speculate that only the normal component of the tensile forces exerted on integrins induces de-adhesion (F ⋅ sin θ, see Fig.7a). De-adhesion then occurs when these tensile forces exceed a threshold set by the strength of integrin adhesion forces. Assuming that the contractile force exerted is constant if adhesion by integrins is experimentally increased or decreased, de-adhesion is expected to occur at a higher or lower angle θ, respectively. To test this hypothesis, we measured the detachment angle θ during wave propagation and compared conditions where we modulated adhesion to the vitelline membrane. *αPS3* RNAi, which reduces adhesion, significantly reduced the detachment angle θ as compared to control embryos. Conversely, *α-cat* RNAi and anterior immobile fences, which we showed increased the duration of the positive feedback and thereby increased integrin adhesion (Fig.6f-g), significantly increased the angle θ relative to controls (Fig.7b-c). Furthermore, depleting *αPS3* in *α-cat* RNAi embryos significantly reduced the angle θ compared to *α-cat* RNAi embryos, similar to *αPS3* RNAi alone (Fig.7b-c). Thus, the angle of detachment changes according to the levels of adhesion to the substrate indicating that tensile forces due to MyoII contractility in the invaginating furrow oppose integrin adhesion forces to the vitelline membrane to induce cortex detachment during wave propagation.

**Figure 4:**
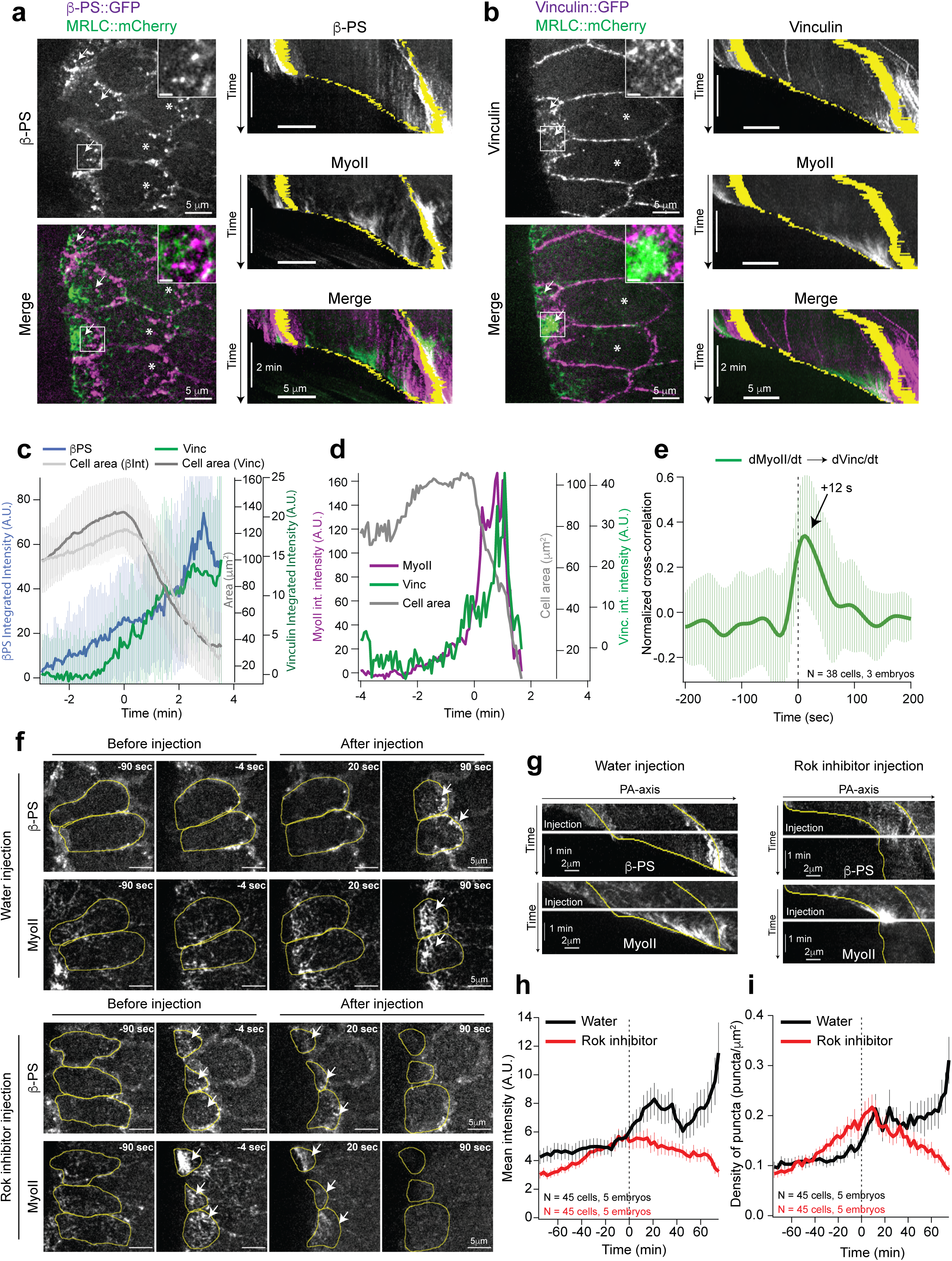
MyoII contractility is required for β-Integrin foci stabilization and maturation. (**a-b**) Representative stills (on the left) and kymographs (on the right) of β-PS (**a**) and Vinculin (**b**) together with MyoII during wave propagation. On the left: grayscale (for β-PS and Vinculin) and merge images. The inset is a zoom on the region within the white box, scalebar 1µm. White arrows indicate cells at the edge of the invaginating furrow where MyoII increases rapidly and propagates as wave. White asterisks indicate cells where MyoII increases slowly and the intracellular wave has not yet been activated. On the right: Greyscale and merged kymographs in representative cells. In yellow the A-and P-boundary of the cell. (**c**) Quantifications of β-PS (in blue) and Vinculin (in green) total integrated intensity over time along with respective quantifications of cell area (in grey). Mean ± SDs between cells. For β-PS N=25 cells from 2 WT embryos, for Vinculin N=42 cells from 3 WT embryos. (**d**) Representative time traces of integrated intensity of MyoII (magenta) and Vinculin (green) along with apical area (grey) in a cell of the propagation zone. (**e**) Average normalized cross-correlation curve between the time derivatives of the MyoII and Vinculin time traces. Mean ± SDs between cells. N=38 cells from 3 WT embryos. (**f**) Stills of β-PS and MyoII in cells of the posterior endoderm from time lapses of water and Rok inhibitor injection during wave propagation. In yellow the cell contours of selected cells. White arrows indicate accumulations of MyoII or integrin focal complexes. (**g**) Representative kymographs along the PA-axis of β-PS and MyoII extracted from cells in **f**. The white line indicates the time of injection. The yellow lines highlight the A-and P-boundaries of one cell. (**h-i**) Quantifications of β-PS mean intensity (**h**) and β-PS density of puncta (**i**) over time in cells at the edge of the invaginating furrow where MyoII increases rapidly. Mean ± s.e.m, N=45 cells from 5 embryos in both water and Rok inhibitor injections. In **d** and **e** time 0 is defined for each cell as the onset of rapid MyoII recruitment (see methods). In **h** and **i** time 0 is the time of injection.

We next studied how fluctuations in adhesion densities in the wild-type impacted the detachment angle θ during wave propagation. Since integrin adhesion is organized in discrete focal complexes where integrins concentrate, the detachment angle θ should fluctuate and increase in front of sites of stronger adhesion (i.e. with more focal complexes). We measured the angle θ over time in WT embryos and found that θ fluctuated during cell detachment, and in particular that θ increased when the local speed of detachment decreased (Fig.7d-e, and Movie 16), as shown by the negative cross-correlation of −0.30±0.02 s.e.m between their time derivatives (Fig.7f). Although the detachment speed was on average similar to the speed of anterior motion of the basal furrow, it fluctuated more (Fig.7d, and Suppl. Fig.7a-d) and transient mismatch between the two speeds highly correlated with the change in angle θ (Suppl. Fig.7e-f). The difference in fluctuations between these two velocities decreased in *α*PS3 RNAi embryos, suggesting that variations in the speed of detachment might be linked to heterogeneities in integrin adhesion to the vitelline membrane. To directly test this, we measured the correlation between the local speed of detachment, measured as the speed of the furrow movement in the most apical sections in contact with the vitelline membrane, to the local βPS intensity and we found that on average the speed of detachment decreased when the local βPS intensity increased (Fig.7g-i).

We showed that the angle of cell detachment from the vitelline membrane changes according to experimentally induced or naturally occurring variations in the strength of integrin adhesion. Together, we conclude that tissue geometry, namely the angle of the furrow at the point of contact with the vitelline membrane, is a tuning parameter that allows cell detachment to occur for a large range of adhesion strength, thereby sustaining wave propagation.

## Discussion

Tissue morphogenesis results from deterministic, pre-patterned information, and self-organized dynamics through feedbacks and local interactions (Collinet and Lecuit, 2021). This is illustrated in the morphogenetic wave of tissue invagination that propagates from the posterior to the anterior in the *Drosophila* embryo (Bailles et al., 2019). Initiation of the wave is determined but its propagation is self-organized. Specifically, the wave is triggered by a genetically controlled invagination in the endoderm primordium (Bailles et al., 2019) and by tissue flow that emerges from the interplay of egg curvature and contractility in the primordium (Gehrels et al., 2023). The tissue-scale wave involves a mechanochemical feedback and interaction with the vitelline membrane (Bailles et al., 2019; Munster et al., 2019), but how the wave emerges from the local interaction between MyoII contractility and adhesion to the vitelline membrane and how cell detachment is regulated was unclear. We now characterized the detailed subcellular mechanisms underlying the tissue scale wave of deformation and reveal a tight interplay between MyoII contractility, integrins adhesion and apical membrane geometry in this process.

We first show that the tissue-scale wave emerges from two mechanochemically coupled subcellular waves: a wave of MyoII contractility within the apical cortex of cells, and a wave of adhesion to and de-adhesion from, the vitelline membrane. Adhesion and MyoII activity are coupled via integrins and depend upon a mechanochemical feedback. De-adhesion is also coupled to adhesion and forward MyoII wave propagation. Thus, adhesion, MyoII activation and de-adhesion form a closed loop that sustains the whole process within cells and across the tissue.

We found that MyoII contractility and cell detachment from the vitelline membrane propagate as coupled waves within cells and not simply from one cell to the next within the tissue, as previously reported (Bailles et al., 2019). Thus, the tissue-scale wave originates from a subcellular mechanochemical process that occurs within the cell cortex and propagates to the next cell as it does within the cortex of a given cell. As such, compartmentalization into cells is not key to understanding what drives wave propagation and a continuum mechanical description of the embryo suffices as previously shown for other processes in the embryo (He et al., 2014). Within the cell we describe two distinct mechanochemical feedback mechanisms that regulate the MyoII wave (Fig.7j). A first mechanism operates at the wavefront in contact with the vitelline membrane and leads to recruitment of new MyoII mini-filaments in an integrin-dependent manner (Fig.3). In contact with the vitelline membrane MyoII contractility also promotes the assembly and maintenance of integrin focal complexes (Fig.4), thus forming a positive feedback loop that amplifies MyoII activation and adhesion. This is similar to the mutual regulation between integrins and actomyosin contractility described in mammals where integrins regulate Rho signaling and MyoII contractility (Guilluy et al., 2011; Schiller and Fassler, 2013; Sun et al., 2016) but also to respond to shear forces exerted by actomyosin contractility (Gauthier and Roca-Cusachs, 2018; Iskratsch et al., 2014; Plotnikov et al., 2012; Raz–Ben Aroush and Wagner, 2006; Schiller and Fassler, 2013). However, it also presents some key differences. Indeed, differently from vertebrates where the Focal Adhesion Kinase (FAK) plays critical roles in integrin-mediated signal transduction and modulates Rho signaling and actomyosin contractility (Guilluy et al., 2011; Schaller, 2010; Tomar and Schlaepfer, 2009), in *Drosophila* FAK null mutants are viable and show only minor defects (Grabbe et al., 2004; Tsai et al., 2008; Ueda et al., 2008). Consistently, we found no defects in MyoII activation and wave propagation in FAK mutants, despite FAK being present in this region of the embryo during gastrulation (data not shown). Thus, in our system integrins activates MyoII via a different mechanism. This might either occur via integrin regulation of other signaling pathways (Sawala et al., 2015b) or it might be mechanical and depend on the coupling between the cortical actomyosin network and integrins (Kovacs et al., 2007). It is interesting to note that this positive feedback depends of the specific configuration of the system in contact with the vitelline membrane where contractile forces due to MyoII are oriented parallel to the substrate and thus exert shear stress on integrin focal complexes. After cortex detachment MyoII further concentrates by being advected with the contracting actin cortex. MyoII induces cortex contraction and promotes its own advection and that of its regulators such as Rho1 and Rok (Munjal et al., 2015). Thus, a second positive feedback operates after cortex detachment that sustains MyoII accumulation while contact with the vitelline membrane by integrins is lost. Interestingly, these two feedback mechanisms are spatially and temporally segregated during wave propagation reflecting sequential adhesion to (first positive feedback) and deadhesion from (second positive feedback) the vitelline membrane (Fig.7j).

Our results further indicate that the kinetics of sequential cell adhesion/de-adhesion to the vitelline membrane has a central role in wave propagation and illustrates the impact of the system’s geometry. First, the contact or not with the vitelline membrane defines whether MyoII locally accumulates by recruitment of new minifilaments or by cortex advection (Fig.7j). Thus, the geometry of the system (in contact or not with the vitelline membrane) instructs the local mechanochemistry of MyoII accumulation. Second, the rate of cell detachment also defines the overall levels of MyoII and adhesion to the substrate by defining the duration of the integrin-MyoII feedback. It is interesting to note that the rate of wave propagation relies on the strength of adhesion to the vitelline membrane. Too much adhesion (as in *α-cat* RNAi and embryos with anterior immobile fences) slows down detachment and brings the system into a vicious cycle: slower detachment further increases adhesion and contractility via the increased duration of the integrin-MyoII feedback, in turn further slowing down wave propagation. Too low adhesion (*αPS3* RNAi) instead fails to sustain the anterior movement of the invaginating furrow and to activate sufficient levels of MyoII. In the wildtype, intermediate levels of integrin adhesion allow smooth anterior propagation of the invagination wave. Third, we report that the tissue geometry offers another level of control through the local angle of detachment and the tensile deadhesion forces (Fig.7l). This sustains wave propagation in spite of fluctuating levels of integrins due to the discrete nature of focal adhesions and of actomyosin asters. Indeed, we showed that the angle of detachment changes according to the levels of adhesion to the substrate in the wild type, as well as in conditions where the levels of adhesion are perturbed significantly. High adhesion as in the case *α-cat* RNAi and embryos with anterior immobile fences increased the angle of detachment while low adhesion as in α*PS3* RNAi embryos decreased this angle (Fig.7l). It should be noted that *α-cat* RNAi may also reduce tensile forces applied to the point of detachment due to impaired force transmission across cells in the invaginated furrow. However, the increased angle is also observed in embryos with anterior immobile fences where force transmission is not affected and where MyoII levels in the invaginated furrow are higher. Altogether, our results illustrate how mechanochemical and geometrical feedbacks integrate locally to regulate cell and tissue dynamics.

Our work also sheds light on new roles of integrin adhesion in morphogenesis. Indeed, integrins are required for the morphogenetic wave in multiple ways. First, integrin-based adhesion initiates the wave through the formation of a deep invagination. The frictional coupling to the vitelline membrane resists the initial anterior flow of the tissue that emerges from tissue curvature(Gehrels et al., 2023) and thereby contributes to the formation of an invagination by tissue buckling. This is important to configure the tissue into a geometry which provides the initial condition for the initiation of the mechanochemical cycle which sustains wave propagation. Second, integrin adhesion also sustains wave propagation. Indeed, integrins amplify the activation of MyoII at the cell cortex and thereby “feed” the invagination with new highly contractile cells that sustain its contraction over time.

The interplay between integrin adhesion and contractility in this morphogenetic process bears interesting similarities with cell motility. Indeed, cell motility and tissue wave morphogenesis in *Drosophila* require both the cyclic interplay between adhesion, MyoII activation and de-adhesion. However, in contrast to cell migration on 2D substrates, in the process we report the retrograde actin flow is not due to actin nucleation at the front of the cells but rather it is retrograde flow of cells (and of their actin cortex) due to a gradient of contractility that peaks in the invagination in the posterior. This is similar to retrograde actin flow in confined 3D cell motility (Liu et al., 2015; Ruprecht et al., 2015) but occurs at the scale of the tissue. In the absence of integrins, posterior contractility in the invagination drives a retrograde flow of cells and their apical actin cortex towards the posterior (Fig.2d) which cannot produce any anterior movement due to the absence of adhesion/friction. In the wildtype, integrin adhesion couples this tissue-level retrograde flow to the vitelline membrane and thereby causes forward movement. Importantly, here the region of integrin-based adhesion to the vitelline membrane propagates also as a wave towards the anterior due to the attachment/detachment cycle we report. If integrin adhesion was irreversible, forward movement would be stalled and wave propagation would be blocked. Sustained forward movement is thus ensured by the fact that adhesion itself is locally transient through cell detachment.

This study exemplifies how large-scale tissue dynamics emerge from the interplay of cell mechanochemical processes and cell geometry.

## Author contribution

C.C., A.B. and T.L. conceived the project and planned experiments. C.C. performed all experiments and quantifications and analyzed the results with T.L. and A.B. A.B. designed the code for the analysis of MyoII rates. C.C. and T.L. wrote the paper and A.B. made comments.

## Acknowledgements

We thank all members of the Lecuit lab for useful feedback during the course of this project. We are particularly grateful to Benoit Dehapiot for help with implementation of KLT and the quantification of MyoII recruitment steps. We are also grateful to Jean-Marc Philippe and Elise Da Silva for generating fly stocks used in this study. We thank the imaging facility at IBDM, member of the France-BioImaging infrastructure supported by the French National Research Agency (ANR-10-INBS-04-01, «Investments for the future»), for assistance with maintenance of the microscope, FlyBase for maintaining curated databases and Bloomington *Drosophila* Stock Center for providing fly stocks. This work was supported by the ERC grant SelfControl #788308. C.C. is supported by the CNRS, T.L. is supported by the Collège de France, A.B. was supported by the ERC grant SelfControl #788308 after a PhD fellowship from the LabEx INFORM (ANR-11-LABX-0054) and of the A∗MIDEX project (ANR-11-IDEX-0001–02), funded by the Investissements d’Avenir French government program.

## Movie legends

**Movie1:** Time lapse of MyoII (MRLC::GFP, greyscale) and Gap-43::mCherry to label cell membranes (in green) in cells of the posterior endoderm during wave propagation in a WT embryo. Note the high level of MyoII activation close to the invaginating furrow and the lower levels more anteriorly. Scalebar 5 µm. Time is in mm:ss.

**Movie 2:** Time lapse of F-Actin (labelled with Utr(ABD)::GFP, greyscale) and MyoII (MRLC::mCherry, green) in a cell of the posterior endoderm at the stage of rapid MyoII recruitment. The yellow arrows indicate discrete MyoII clusters appearing just prior cell detachment. Scalebar 5 µm. Time is in mm:ss.

**Movie 3:** Top and side views of F-actin (labelled with Utr(ABD)::GFP) of a cell during apical cortex spreading onto the vitelline membrane and wave propagation. The cell contours are in yellow. The yellow vertical lines in the side view indicate the A-and P-boundaries of the cell. Scalebars: 5 µm in the top view and 2 µm in the side view. Time is in mm:ss.

**Movie 4:** Time lapse of β-PS::GFP in cells of the posterior endoderm during wave propagation. The white arrows point to β-PS puncta in the medio-apical cortex of cells. Scalebar 5 µm. Time is in mm:ss.

**Movie 5:** Time lapse of Talin-I-Ypet in cells of the posterior endoderm during wave propagation. The white arrows point to Talin puncta in the medio-apical cortex of cells. Scalebar 5 µm. Time is in mm:ss.

**Movie 6:** Top and side views MyoII (MRLC::mCherry) propagation in a cell of the posterior endoderm. The cell contours are in yellow. The white arrows indicate the onset of MyoII recruitment in clusters which occurs in contact with the vitelline membrane. The cell contours are in yellow. Scalebars are 5 µm for both the top and side view. Time is in mm:ss.

**Movie 7:** Time lapse of E-cad (E-cad::GFP, in magenta) and MyoII (MRLC::mCherry, in green) in a control (water injection, top) and an αPS3 RNAi injected embryo. Scalebar 10 µm. Time is in mm:ss.

**Movie 8:** Time lapse of β-PS (β-PS::GFP, in magenta) and MyoII (MRLC::mCherry, in green) in cells of the posterior endoderm during wave propagation in a WT embryo. The merge view is on the right and the corresponding β-PS signal in greyscale is on the left. The white arrows point to β-PS puncta in the medio-apical cortex of cells. Scalebar 5 µm. Time is in mm:ss.

**Movie 9:** Time lapse of Vinculin (sGFP::Vinculin, in magenta) and MyoII (MRLC::mCherry, in green) in cells of the posterior endoderm during wave propagation in a WT embryo. The merge view is on the right and the corresponding Vinculin signal in greyscale is on the left. The white arrows point to Vinculin puncta in the medio-apical cortex of cells. Scalebar 5 µm. Time is in mm:ss.

**Movie 10:** Time-lapse of β-PS (β-PS::GFP, on the top) and MyoII (MRLC::mCherry, at the bottom) in a water (left) or Rok inhibitor (H-1152) injection during wave propagation. The white arrows indicate formed β-PS focal complexes at the time of injection. Scalebar 5 µm. Time is in mm:ss.

**Movie 11:** Tissue-level side views of a WT embryo (Control) and an embryo with an anterior immobile fence (AF). E-cad (E-cad::GFP) is in magenta and MyoII (MRLC::mCherry) is in green. Scalebar 20 µm. Time is in mm:ss.

**Movie 12:** Time lapse of MyoII (MRLC::mCherry) during wave propagation in a control (water injection, top) an αCat RNAi (middle) and an embryo with an anterior immobile fence (AF, bottom). Scalebar 20 µm. Time is in mm:ss.

**Movie 13:** Time lapse of MyoII (MRLC::mCherry) during wave propagation in a control (water injection, top left), an αCat RNAi (top right), αPS3 RNAi (bottom left) and an embryo injected with both RNAi against αCat and αPS3 (αCat + αPS3 RNAi, bottom right). Scalebar 5 µm. Time is in mm:ss.

**Movie 14:** Time lapse of β-PS (β-PS::GFP, in greyscale) and MyoII (MRLC::mCherry, in green) in cells of the posterior endoderm during wave propagation in a control (water injection) embryo (on the top) and an embryo injected with αCat RNAi (at the bottom). The merge view is on the left and the corresponding β-PS signal in greyscale is on the right. Scalebar 5 µm. Time is in mm:ss.

**Movie 15:** Time lapse of β-PS (β-PS::GFP, in greyscale) and MyoII (MRLC::mCherry, in green) in cells of the posterior endoderm during wave propagation in a WT embryo (on the top) and an embryo injected with an anterior immobile fence (at the bottom). The merge view is on the left and the corresponding β-PS signal in greyscale is on the right. Scalebar 5 µm. Time is in mm:ss.

**Movie 16:** Side view of a cell detaching from the vitelline membrane during wave propagation in a WT embryo. F-Actin is labelled with Utr(ABD)::GFP. Scalebar 5 µm. Time is in mm:ss.

## STAR Methods

### Fly strains

Fly lines carrying following insertions were used: live cell imaging of MyoII Regulatory Light Chain (MRLC) (encoded by the gene *spaghetti-squash*, *sqh,* FlyBase ID: FBgn0003514) was carried out using either *sqh-MRLC::mCherry* (either inserted on chromosome 2 at the VK18 [53B2] site (Bailles et al., 2019) or inserted at the VK27 [89E11] site (Garcia De Las Bayonas et al., 2019) on chromosome 3) or *sqh-MRLC::EGFP* on chromosome 2 (gift from R. Karess). F-Actin was visualized using a *sqh-Utr(ABD)::GFP* insertion on chromosome 3 (Dehapiot et al., 2020; Rauzi et al., 2010). Ubiquitous expression of a *UASp-Gap43::mCherry* insertion on chromosome 3 was used to visualize the plasma membrane (Guillot and Lecuit, 2013) and was driven using a *nanos-Gal4*. E-cadherin (*shg* in Drosophila, FlyBase ID: FBgn0003391) was visualized using *E-cad::EGFP^Kin^*, an EGFP knock-in allele at the locus generated by homologous recombination (Huang et al., 2009). β-PS is the main β-Integrin in *Drosophila* and it is encoded by the gene *myospheroid* (*mys*, FlyBase ID: FBgn0004657). Live imaging of β-PS was carried out using *mys::EGFP^Kin^* an EGFP knock-in allele of *myospheroid* at the locus (Klapholz et al., 2015). Talin is encoded by *rhea* (FlyBase ID: FBgn0260442) in *Drosophila* and a Y-Pet knock-in allele in exon 6 (Talin-I-Ypet) (Lemke et al., 2019) was used for live imaging. Vinculin vas visualized using an *sGFP::Vinculin* fosmid insertion on Chromosome 3 at the ATTp-2 landing site (Kale et al., 2018). Rho1-GTP was visualized using *ubi-Anilin(RBD)::EGFP* which functions as Rho1 activity sensor (Munjal et al., 2015).

All used fly strains are listed in Supplementary Table 1.

### RNA interference and drug injections

dsRNA probes directed against αPS3 (*scb*, CG8095, FlyBase ID: FBgn0286785) and α-Catenin (*aCat*, CG17947, FlyBase ID: FBgn0010215) were prepared and injected at a final concentration of 5 μM in embryos less than 1 h old as previously described(Bailles et al., 2019; Cavey et al., 2008). The Rho kinase (Rok in *Drosophila*, FlyBase ID: FBgn0026181) inhibitor H-1152 (Enzo Lifesciences) was injected at a concentration of 40 mM in embryos at stage 7 during time lapse using an InjectMan4 micromanipulator and a FemtoJet 4i microinjector from Eppendorf directly installed on the microscope.

The dilution factor in the embryo is typically 1:50 so we expect a final concentration of 800 μM for the H-1152 inhibitor and 100 nM for the dsRNA probes.

Injections were performed at 50–80% embryo length from the posterior and on the lateral side in all experiments in this paper.

The sequences of the primers used to generate the dsRNA probes against α*PS3* and *αCat* are available in (Bailles et al., 2019) and (Cavey et al., 2008) respectively.

### Live Imaging

For live imaging embryos were prepared as described earlier (Cavey and Lecuit, 2008) and imaged at stage 7 in the dorsal and posterior region (unless otherwise specified) for 12 to 15 min at room temperature (22°C) depending on the experiment. Dual color time lapse imaging was performed using simultaneous acquisition on a confocal spinning disc (CSU-X1, Yokogawa) Nikon Eclipse Ti inverted microscope equipped with 2 cameras (Rolera EM-C^2^, QImaging) using a 100X/N.A 1.45 Plan Apo oil-immersion or a 40X/1.25 NA Apo water-immersion objectives from Nikon.

For live imaging of F-Actin and MyoII z-stacks of 4 µm (8 planes, spacing 0.5 µm) from the vitelline membrane were acquired with a time interval 4s/stack. For live imaging of β-PS alone or together with MyoII or Vinculin and of Talin z-stacks of 1 µm (2 planes, spacing 0.5 µm) from the vitelline membrane were acquired with a time interval 3s/stack. For live imaging of Vinculin and MyoII z-stacks of 2 µm (4 planes, spacing 0.5 µm) from the vitelline membrane were acquired with a time interval 4s/stack. For live imaging of E-cad and MyoII (Fig.3e) z-stacks of 6 µm (12 planes, spacing 0.5 µm) from the vitelline membrane were acquired with a time interval 6s/stack. Time lapses of deeper stacks (10 µm) of MyoII (Fig.3a) were acquired by collecting 20 planes with a spacing of 0.5 µm every 10s. Deep z-stacks (9-10 µm) of β-PS and Talin (Suppl. Fig. 2b-c) were acquired in living embryos with a spacing of 0.5 µm. The stacks form where side views of E-cad and MyoII in Movie 11 were extracted were acquired by collecting 33 planes with a spacing of 0.8 µm every 20s.

In all cases, imaging conditions (exposure time and laser power) were optimized and kept constant between controls and perturbed embryos.

### Generation of tissue fences by tissue cauterizations

Tissue cauterizations to perturb the anterior movement of the dorsal epithelium were generated as previously described (Bailles et al., 2019). Briefly, a line cauterization of approximately 30 µm along the Medio-Lateral (ML) axis at ∼30% Embryo Length (EL) from the posterior was generated by focusing a near-infrared (NIR, 1030nm) laser with a 100X/N.A. 1.4 oil-immersion Plan Apo VC, Nikon objective on the apical side of the embryonic blastoderm (∼1–2 μm above adherens junctions) with an average power of 500 mW at the back aperture of the objective and moving the stage with a constant speed of 17-20 μm/s.

### Image processing, and data analysis

#### Software

All image processing and data analysis were performed using ImageJ (1.53v), Matlab 2019b including Curve Fitting, Image Processing, Statistics and Machine Learning Toolboxes (Mathworks), Ilastik (1.3.3) and IgorPro (v6.3 or 9, Wavemetrics). Graphs were produced with either Matlab or IgorPro and exported to Adobe Illustrator 2023 for final processing in figures.

#### Image processing, cell tracking and segmentation

Maximum intensity projections were used for measurements of fluorescence intensity and for cell segmentation. Before projection z-stacks were smoothened with a mean filter with a kernel of 0.04 µm (0.5 pixels) to increase the signal-to-noise ratio (SNR) in the projected images. Cell segmentation and tracking were performed as previously described (Collinet et al., 2015) using Ilastik (1.4.0. rc2) and Tissue Analyzer (Aigouy et al., 2010).

Side views along the AP-axis were generated by first registering cells along the y direction using the position of cell centroids to remove y drifts and by re-slicing the image stacks across a DV width of 2 µm (25 pixels) around the cell centroid. The side views are then a maximum intensity projection of the resliced planes. Kymographs along the AP-axis were obtained with a similar method but using maximum intensity projection time series as input for re-slicing. Kymographs are either maximum intensity projections across a DV width of 2 µm around the cell centroid (Figs 1e, 2g-h, 4a-b, and 4g) or across the entire medio-apical region of the cell (Fig. 5b).

**Figure 5:**
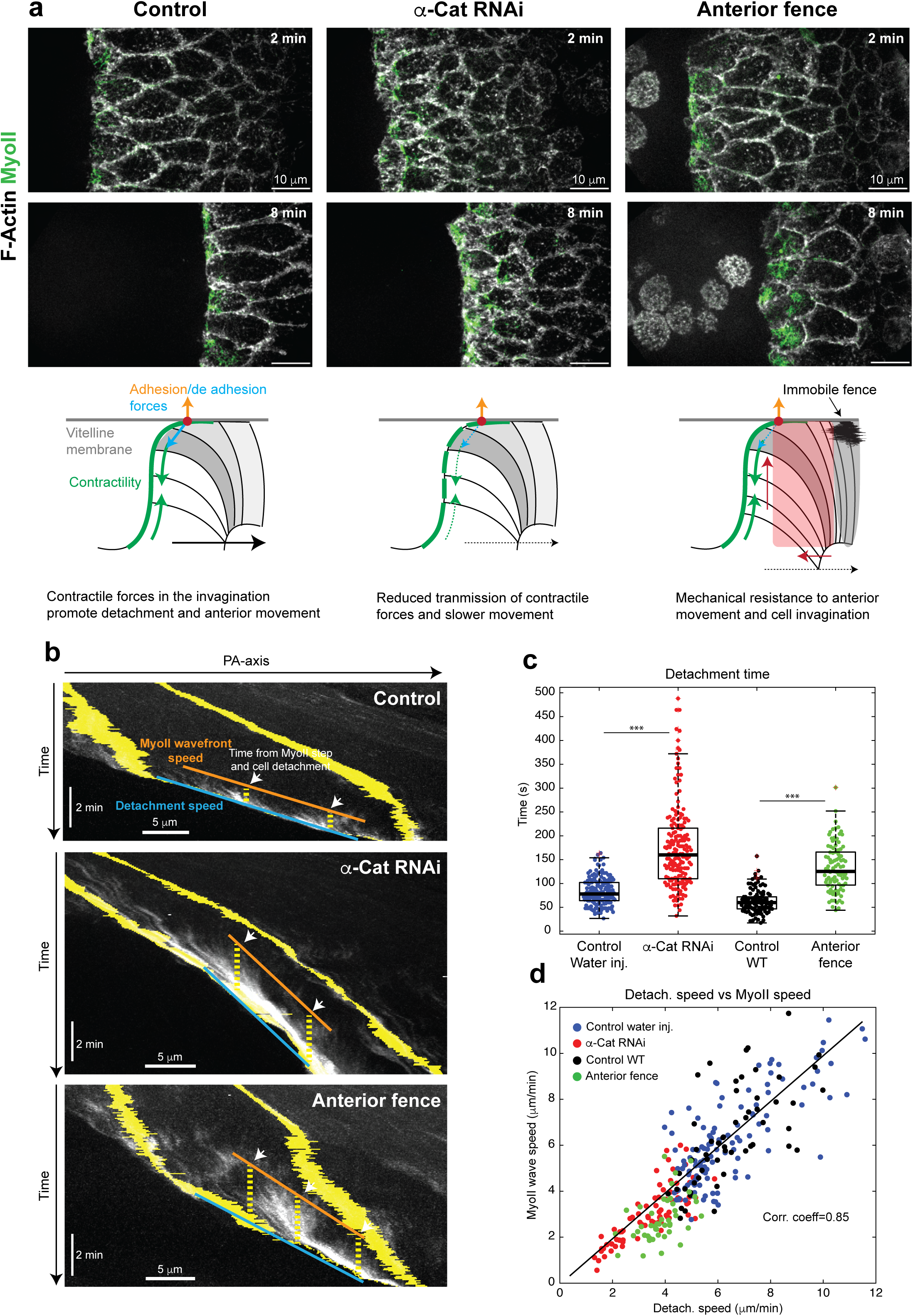
Reducing detachment speed reduces MyoII wavefront speed. (**a**) Top: High resolution tissue-level view of MyoII propagation in a WT (left), an α-Catenin RNAi (middle) and an embryo with anterior immobile fences embryo (right). F-Actin is in greyscale and MyoII in green. Bottom: Cartoon of the forces involved in cortex detachment of the cell at the front of the invaginating furrow in the corresponding conditions above. Green arrows: contractile forces, orange and blue arrows: adhesion and de-adhesion forces, the black arrows indicate the anterior movement, the red arrows indicate the resistive forces due to the immobile fences and the red circle is the point of cortex detachment. Dashed lines indicate reduction of the speed of movement (black arrows) or force integration (green arrows) compared to WT. (**b**) Kymographs along the PA-axis of MyoII in a control (WT), an α-Catenin depleted cell and a cell from an embryo with anterior fences. In yellow: the A-and P-boundaries of the cell. White arrows: MyoII accumulations. The orange and cyan lines indicate the speed of the MyoII wavefront and the speed of cortex detachment respectively. The time of local cortex detachment following MyoII accumulation is labelled by the yellow dashed line. (**c**) Measurements of local cortex detachment time following MyoII cluster accumulation. Individual data points are superimposed to box plots. N=161 events in 95 cells from 5 embryos for water injection, 185 events in 89 cells from 5 embryos for α-Cat RNAi, 113 events 57 cells from 3 embryos for WT control and 99 events in 45 cells from 3 embryos for Anterior fence. *** indicates a p-value <0.001 at a Mann-Whitney test. (**d**) Scatter plot of the detachment vs MyoII wavefront speed of propagation measured as illustrated in **b**. N=116 cells from 5 embryos for water injection, 75 cells from 5 embryos for α-Cat RNAi, 57 cells from 3 embryos for WT control and 45 cells from 3 embryos for Anterior fence. The pearson correlation coefficient is 0.85 with a p<0.001.

The apico-basal position of the actin cortex was estimated using a custom ImageJ macro integrating the ‘stack focuser’ plugin from M. Umorin previously described (Bailles et al., 2019; Collinet et al., 2015). The macro uses the ‘stack focuser’ plugin to define the most focused plane in all positions of the image. This information is stored in an heightmap image which is smoothened with a median filter using a radius of 1.2 µm (15 pixels). The smoothened images are then used as input to generate heatmap kymographs and spatial profiles of the apico-basal position of the cortex as described below. All measurements of fluorescence were performed after local background subtraction performed as previously described(Bailles et al., 2019; Collinet et al., 2015). Briefly, for all images background intensities were subtracted by masking out structures of interest (MyoII clusters, Integrin puncta etc.) and then subtracting residual intensities. The masks were generated using a manually defined intensity threshold on images pre-treated with the “background subtraction” function of ImageJ (using a radius of ∼4 μm (50 pixels) for MyoII and ∼0.4 μm (5 pixels) for βPS and Vinculin) to reduce the effects of uneven illumination of view field. Residual background images were smoothened with a gaussian blur filter with a radius of ∼0.3-0.6 μm (4-8 pixels) before subtraction.

#### Measurements of fluorescent intensity and of focal complex puncta parameters in cells

Measurements of fluorescence intensity were performed in the entire medio-apical region of cells (Figs. 1c, 2i-l, 4c-d, 4h-i, 6d and Suppl Figs. 1a, 3a-d, 5c-d) using an ROI obtained by automated segmentation and tracking that was shrunken by 10 pixels (0.8 µm) to remove junctional signal.

For measurements of focal complex puncta, a simple threshold-based image segmentation was performed to detect individual puncta and extract their size and intensity. First, a manually defined threshold was selected on images pre-treated with the “background subtraction” function of ImageJ (using a radius of ∼0.4 μm) to reduce the effects of uneven illumination of view field. Then images were binarized and particles were detected with the function “analyze particles” of ImageJ. The particles belonging to each cell were used to determine the total number of puncta per cell, their average size, mean intensity and integrated intensity. In Fig. 4i, Suppl. Fig. 5d and Fig.6g the density of puncta per cell was determined as N of puncta per cell / cell area. For β-PS a running average across 3 images of the time lapse was performed before particle detection to further reduce noise and fluctuations in particle detection.

Time registration of individual cells was always relative to the onset of the rapid MyoII recruitment phase and was performed on measurements of MyoII mean intensity across the entire cell. Time 0 was initially defined when MyoII mean intensity overcame a threshold and then was corrected manually based on cell area and MyoII integrated intensity measurements (this is also the time when the area of the cell starts decreasing due to its detachment from the vitelline membrane). The same time registration was used to register cells for the generation of heatmap kymographs (see below).

#### KLT feature tracking, MyoII rates estimation and heatmap kymographs genetation

##### Measurement of the velocity field

Velocity fields of the F-actin network were obtained by tracking discrete apical F-actin structures with a Kanade-Lucas-Tomasi (KLT) (Lucas and Kanade, 1981; Tomasi and Detection, 1991) feature tracking algorithm coupled with an affine consistency test to minimize errors (Shi, 1994) implemented in the C programming language (Stan Birchfield https://cecas.clemson.edu/~stb/klt) as previously described (Dehapiot et al., 2020). The KLT output is then post-processed using a custom Matlab routine. Trajectories were filtered to eliminate too short trajectories (less than 3 frames, i.e. 8 sec), those with unrealistic particle speeds (more than 0.24 μm/sec) and those crossing boundaries between two cells. From the particles trajectories, a velocity field is obtained by derivating the position of all the particles and interpolating the resulting velocities on a fixed grid of 0.64×0.64μm^2^ (8×8 pixels^2^). We compared results obtained with KLT to Particle Image Velocimetry (PIV) and found that KLT is computationally faster and gives comparable results. In our case, KLT, since it tracks features, has the advantage that the velocity field is not influenced by material coming into the field of view from other planes (reviewed also in (Vig et al., 2016)).

##### MyoII mass balance in the plane of the cortex

The evolution of MyoII concentration in a given volume of the cortex, here considered a 2D surface, is set by the MyoII bound to the actin cortex and being transported with it (by advection within the plane), and by the net balance of binding/unbinding of new MyoII minifilaments to the cortex (i.e. from the cytoplasmic pool). As previously described for actin concentration (Vallotton et al., 2004) and MyoII (Nishikawa et al., 2017), we use the mass balance equation:

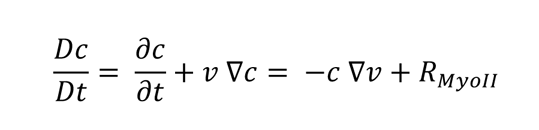

Where *c* is MyoII concentration, *v* the velocity of the MyoII flow, and *R_Myoll_* is the effective rate of Myosin incorporation (net binding/unbinding rate of new MyoII minifilaments) into the actin cortex, and *DC/Dt* is the comoving derivative of the concentration. We infer the speed of MyoII within the plane %’ from the velocity field of actin particles (see above) and the concentration of MyoII from the fluorescence intensity of MRLC::mCherry in the image (maximum intensity projections obtained as described above). Pixel intensities are averaged on a grid of 0.64×0.64μm^2^ (8×8 pixels^2^), then a rolling average on time is performed, with a window of 3 frames (8 seconds). We then calculate spatial and temporal derivatives using respectively central differences and forward difference. We can thus deduce from the previous equation the effective reaction kinetics rate (R_MyoII_) for each position of the grid at all time points:

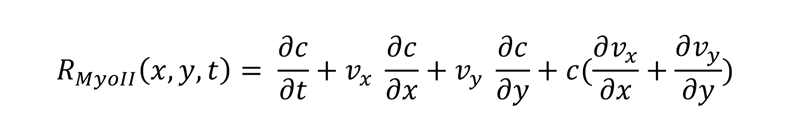

We then average *R_Myoll_* in time with a rolling average of 3 frames (8 seconds).

The validity of our code was checked on simple 1D simulations where we impose a homogeneous expansion rate of the cells, and keep MyoII intensity constant. The average R_MyoII_ value found by our method described above, although noisy at the boundary of the MyoII domain, matched the expected value of R_MyoII_.

Since we are interested in the time evolution of MyoII bound to a region of the actin cortex which can move by translations, in Figs 3c, 3g, 6b, 6e and Suppl. Figs. 4e and 6b we plotted the comoving time derivative of MyoII *DC/Dt*. Indeed, being our analysis based on an Eulerian representation (fixed grid), the movement of the actin cortex between two arbitrarily defined neighboring boxes (e.g. from box 1 to box 2) causes corresponding changes of the concentration time derivative *DC/Dt* between the)’boxes (e.g. an increase in box 2 and a corresponding decrease in box 1). The term *v*Δ*c* accounts for this, such that comoving time derivative *DC/Dt* is not affected by simple movements between neighboring %’ boxes, but only by concentration by advection with the contracting cortex (−*c* ∇*v*) and net binding/unbinding rate of new MyoII minifilaments to the cortex (*R_Myoll_*).

##### Generation of the reconstructed kymographs heatmaps

At each time point we attribute each box of the grid (see paragraph above) to a given cell using the information from cell segmentation and tracking. The position of the boxes is then registered together with the cell. In time, cells are registered as described above using measurements of MyoII mean intensity in the entire cell. In space, cells are registered either by positioning their P-boundary at time 0 at the position x=0 as shown in Suppl Fig. 1d or by defining x=0 the position of the cell P-boundary for all time points (moving P-boundary), as in Suppl. Fig. 1e. We then average (by the mean, except for the deformation rate where an averaging by the median was performed) in space (along the dorsoventral axis) and across different cells, to obtain the kymograph as in Figs 1g, 3f and Suppl. Figs. 1d-e, 4a-c.

Spatial and temporal profiles of the MyoII concentration, of the MyoII rates, of cell deformation and of the cortex apico-basal position were obtained for each embryo directly from the heatmap kymographs by computing the median across the y-axis (for spatial profiles) or the mean across the x-axis (for temporal profiles (as illustrated in Suppl. Fig. 4a and 4c). For spatial profiles, heatmap kymographs in the frame of reference of the cell moving P-boundaries were used. For temporal profiles, we used heatmap kymographs in the fixed frame of reference of the embryo (cell registration relative to position of the P-boundary at time 0). The spatial profile of each embryo was obtained by computing the median the between 0-3 min (or 0-6 min for *acat* and *acat + αPS3* RNAi in Fig.6e) at all x positions in the heatmap. The temporal profile of each embryo was obtained by computing the mean across a region of 5 µm (centered around x=4-7 µm depending on the embryo) for all y positions (time) in the heatmap. Profiles of the apico-basal position of the cortex were obtained with the same method using heightmap images obtained as described above.

Heatmap kymographs of MyoII concentration in the fixed frame of reference of the embryo were used to estimate the speed of the MyoII wave inside cells in Fig.1g. Similar to what previously described (Bailles et al., 2019), the mean heatmap kymograph across 3 embryos was first binarized and then a linear fit of the positions where MyoII concentration was higher than a threshold was performed as a function of time. Note that the speed extracted here was compared to the speed of the wave in fixed frame of reference of the embryo as in Ext. Fig. 6j in (Bailles et al., 2019) and not the frame of reference of the tissue as in Fig.1d of the same paper. In this latter reference a correction for the elastic deformation of the cells due to their apical spreading to the vitelline membrane was applied.

#### Detection of steps of the MyoII wave, measurements of MyoII wavefront speed, detachment speed and detachment time from kymographs

The discrete events of MyoII recruitment during the propagation of the intracellular MyoII wave were detected manually by comparing for each cell top views (max projections) of MyoII with corresponding time traces of the integrated fluorescence intensity. Steps were detected when significant increases in the MyoII integrated fluorescence intensity were associated to visible recruitment of MyoII in the medio-apical region of the cell. The number of MyoII recruitment steps after the max in cell area (i.e. during intercellular wave propagation) is reported in Fig.1f.

The measurements of the MyoII wavefront and detachment speed in Fig.5b were performed on kymographs of MyoII using the Velocity Measurement Tool (available at https://dev.mri.cnrs.fr/projects/imagej-macros/wiki/Velocity_Measurement_Tool). Kymographs of MyoII were produced as described above. For each cell, steps of MyoII accumulations detected as above were reported in the corresponding MyoII kymographs. A segmented line connecting the anterior front of such clusters accumulations was drawn manually and the average MyoII wavefront velocity was computed using the function “measure Velocities” of the Velocity Measurement Tool. The detachment speed was computed similarly by manually drawing a segmented line following the detaching P-boundary of the cell. The time of detachment was computed by measuring the y distance (in seconds) between the two segmented lines at center of MyoII clusters accumulations.

#### Measurements of the angles θ and of detachment point and furrow velocities from side views

The detachment angle θ, and the velocities of the detachment point and of the basal furrow were measured from reconstructed side views of F-actin obtained as described above. The detachment point was defined as the most posterior point of the cell still in contact with the vitelline membrane. The position of the basal furrow was defined as the most basal point at the posterior of the cell visible in our stacks. The position of these points was tracked manually along the AP-axis and instant velocities were computed. The detachment angle θ at all time points was determined from the coordinates of these two points using the atan2 (arctangent) function with the ImageJ software.

#### Measurements of β-PS intensity in front of the advancing furrow and correlation with the detachment speed

To correlate the local intensity of β-PS to the local detachment speed (Fig.7h-i), we manually tracked for each embryo the point of detachment (defined as the most posterior point of the tissue anterior to the furrow) along 4-5 horizontal lines (yellow line Fig.7g) along the AP-axis and we measured the β-PS intensity just anterior to this point. The instantaneous AP detachment velocity was calculated from the x coordinates of the tracked points. The local β-PS intensity mean intensity was measured in an oval region with an aspect ratio of 0.66 and a major DV axis of 6 µm (75 pixels) positioned 0.4 µm (5 pixels) anterior to the point of detachment for each detachment point and for all time points.

**Figure 6:**
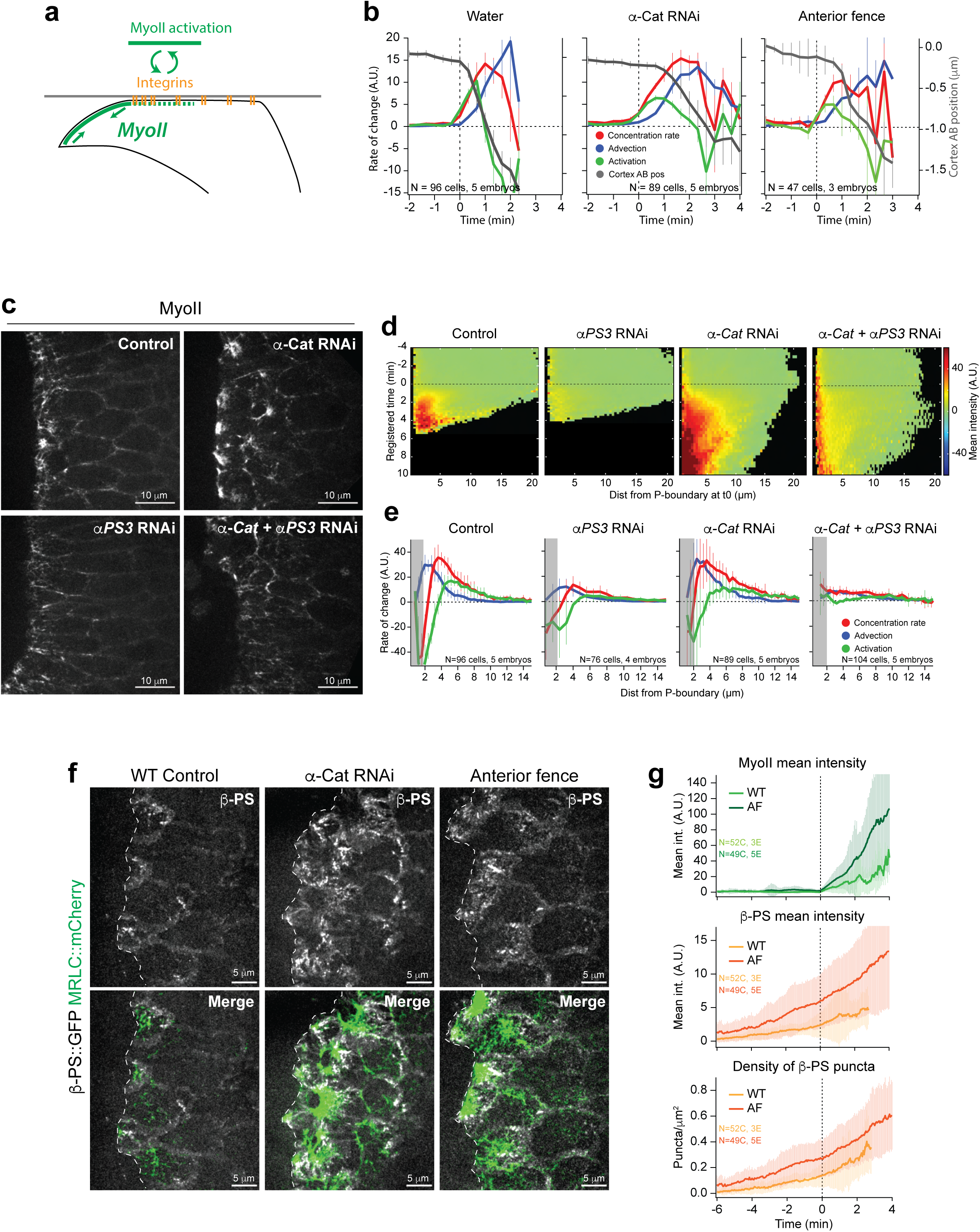
Prolonging contact with the vitelline membrane increases the duration of the Integrin-MyoII feedback. (**a**) Cartoon illustration of the positive integrin-MyoII feedback in contact with the vitelline membrane. (**b**) Time evolution of the MyoII rates along with the cortex apico-basal position in a fixed region of the cortex in the indicated conditions. N= 5 embryos (96 cells) for water injection, 5 embryos (89 cells) for α-Cat RNAi and 3 embryos (47 cells) for anterior fence. Mean ± s.e.m between embryos. (**c**) Stills from time lapses of MyoII in the indicated conditions. Note the reduction of MyoII levels in *α-Cat+αPS3* RNAi embryos compared to *α-Cat* RNAi alone. The MyoII levels are similar to *αPS3* RNAi alone. (**d**) Heatmap kymographs of MyoII mean intensity in the indicated conditions. (**e**) Average spatial profile of MyoII rates in the indicated conditions (averaging time window 0-3min for WT and *αPS3* RNAi and 0-6 min for *α-Cat* and *α-Cat+αPS3* RNAi). The gray boxes indicate where cells disappear from the field of view and measurements become less reliable. In both **d** and **e** the data are plotted relative to the moving P-boundary of the cells and N=96 cells from 5 control (water injected), 76 cells from 4 *αPS3* RNAi, 89 cells from 5 *α-Cat* RNAi and 104 cells from 5 *α-Cat+ αPS3* RNAi embryos. In **e** mean ± S.D. between embryos are shown. (**f**) Representative stills from time lapses of β-PS and MyoII in the indicated conditions. Grayscale for β-PS and merge images are shown. The dashed white line indicates the border of the advancing invaginating furrow. (**g**) Time traces of MyoII mean intensity (top), β-PS mean intensity (middle) and the density of β-PS puncta (bottom) in WT (light green and light orange) and embryos with anterior fences (AF, dark green and dark orange). N=52 cells from 3 WT and 49 cells from 5 embryos with anterior fences. Mean ± SDs between cells. In **b**, **d** and **g** time 0 is the time of wave activation (rapid MyoII recruitment, see methods).

**Figure 7:**
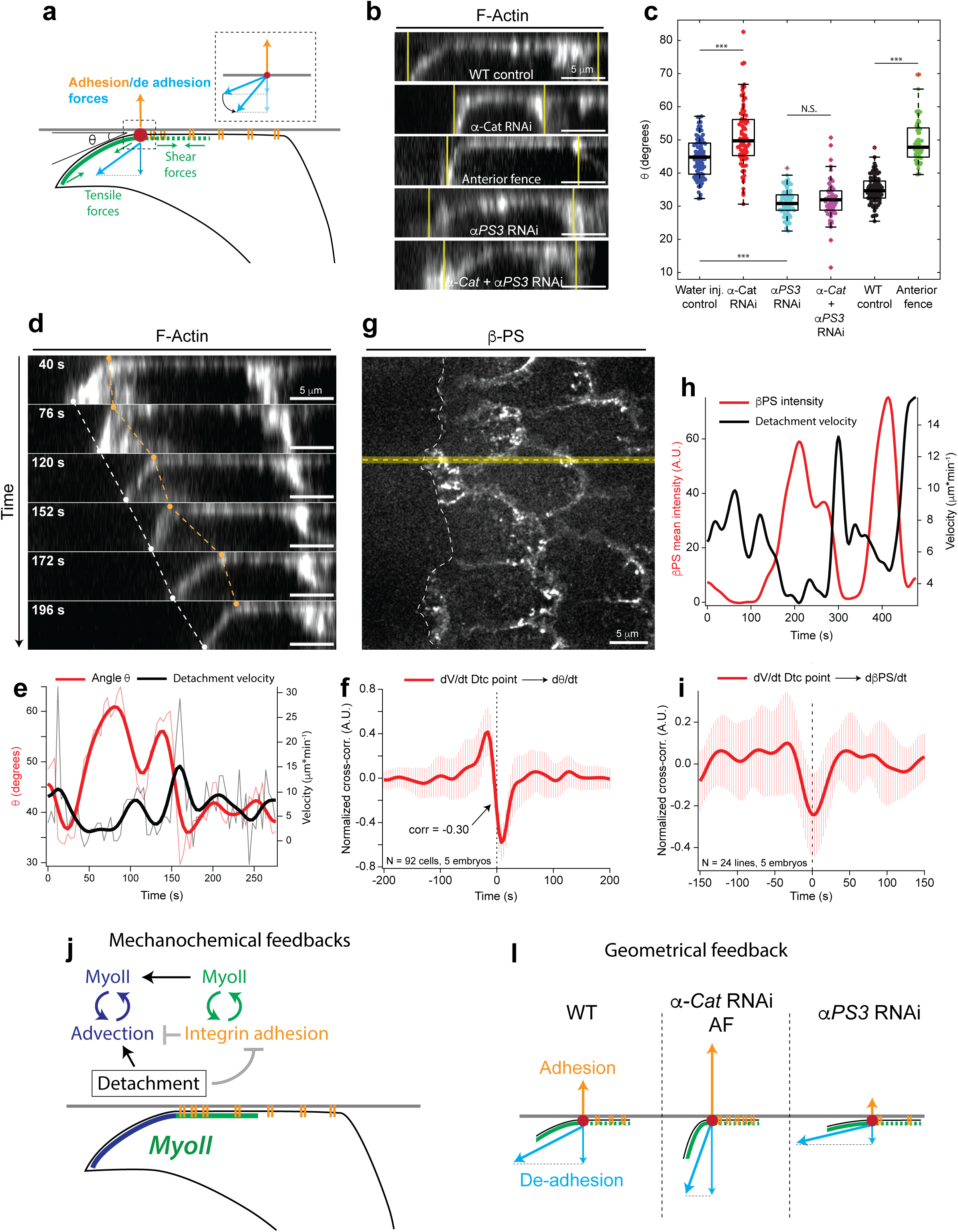
The local angle of detachment changes according to the adhesion strength. (**a**) Illustration of the forces acting at the site of detachment in cells during wave propagation. Shear and tensile forces in integrin complexes due to MyoII contractility (in green) are represented. MyoII contractility in the invaginated furrow and the detached side of the cell gives rise to a de-adhesion force (in blue) at the site of detachment (red circle) opposing adhesion (in orange). The component of the de-adhesion force on integrins in the direction perpendicular to the plane of adhesion depends on the angle θ of detachment. The inset illustrates how increasing the angle θ increases this component for a given de-adhesion force. (**b**) Representative side views of cells during wave propagation in the indicated conditions. The yellow vertical lines indicate the A-and P-boundaries of the cells as detected from cell segmentation of corresponding top views. (**c**) Quantifications of the average detachment angle θ in the indicated conditions. Individual data points are superimposed to box plots. N=95 cells from 5 embryos for Water injection control, 85 cells from 5 embryos for α-Cat RNAi, 76 cell from 4 embryos for *αPS3* RNAi, 60 cells from 5 embryos for *α-Cat+ αPS3* RNAi, 98 cells from 6 embryos for WT control and 47 cells from 3 embryos for anterior fence. *** indicates a p-value <0.001 and N.S. a p-value>0.05 at a Mann-Whitney test. (**d**) Side views from a time lapse of cell detachment in a WT embryo. The white dots and the white dashed line indicate the movement of the invaginated furrow (as basal as it can be detected in our stacks). The orange dots and the orange dashed line indicate the movement of the detachment point. (**e**) Time traces of the angle θ (in red) and the speed of the detachment point along the PA axis (in black) of the cell in **d**. The thick lines are smoothened time traces and the thin semi-transparent lines are the actual measurements from time lapses. (**f**) Average normalized cross-correlation curve between the time derivatives of the detachment speed and the angle θ time traces. Mean ± SDs between cells. N=92 cells from 5 WT embryos. (**g**) Apical view of the detachment front of the invagination in a WT embryo labelled with β-PS::GFP. The white dashed line labels the detachment front. The yellow horizontal line is an example of horizontal lines used to measure the speed of the detachment front and the local β-PS intensity (see methods). (**h**) Representative smoothened time traces of the local β-PS mean intensity (in red) and the detachment front velocity along the PA-axis (in black) measured along an horizontal line as that in yellow in **g**. (**i**) Average normalized cross-correlation curve between the time derivatives of the detachment point speed and the local β-PS intensity time traces. Mean ± SDs between cells. N=24 lines from 5 WT embryos. (**j**) Cartoon illustration of the mechanochemical feedbacks recruiting MyoII during wave propagation. At the wavefront the integrin-MyoII positive feedback locally activates MyoII and reinforces adhesion. After cortex detachment MyoII is further recruited by advection with the contracting cortex. Integrin adhesion blocks cortex contraction and detachment interrupts the integrin-MyoII positive feedback and thus promotes advection. (**l**) Cartoon illustration of the change in the detachment angle θ according to different levels of adhesion force. Examples of the conditions showing these effects are listed.

#### Cross-correlation

Cross-correlation was performed as previously described(Collinet et al., 2015). Briefly, time traces were smoothened using a binomial algorithm using IgorPro software before derivation. The normalized cross-correlation was calculated with IgorPro software using the following formula:

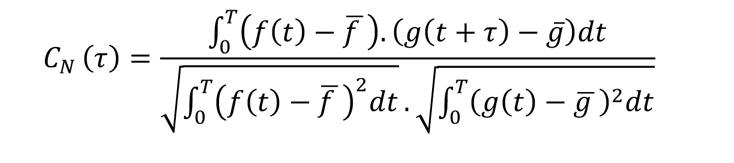

where *t* is the time, *T* the total time of analysis, *τ* the lagtime. The mean cross-correlation function was calculated as the average of the cross-correlation functions of each cell or line (Fig.7i) and was used to estimate the peak correlation value and the corresponding time delay. The frame rate of the time lapses used for this analysis was 3 or 4 s/frame depending on the experiment.

### Statistics

For all experiments data points from different steps/lines/cells/embryos from at least 2 independent experiments were pooled to estimate the mean, S.D. and s.e.m. values plotted. All of the *P* values are calculated using a non-parametric Mann–Whitney test (Matlab statistic toolbox). No statistical method was used to predetermine sample size. The experiments were not randomized, and the investigators were not blinded to allocation during experiments and outcome assessment.

### Repeatability

All measurements were performed in 2-5 embryos. In experiments where embryos are not injected, we consider each embryo as an independent experiment. In drug and dsRNA injection experiments the number of independent experiments is defined as the number of independent injections. Representative images, which are shown in Figs 1-7 and Suppl. Figs 1-7 were repeated at least twice and up to more than ten times.

### Data availability

All the data supporting the findings of this study are available within the paper. Raw image data are available upon reasonable request.

### Code availability

The custom codes used to process images analyze data are available upon request.

---

**Supplementary Figure 1:**
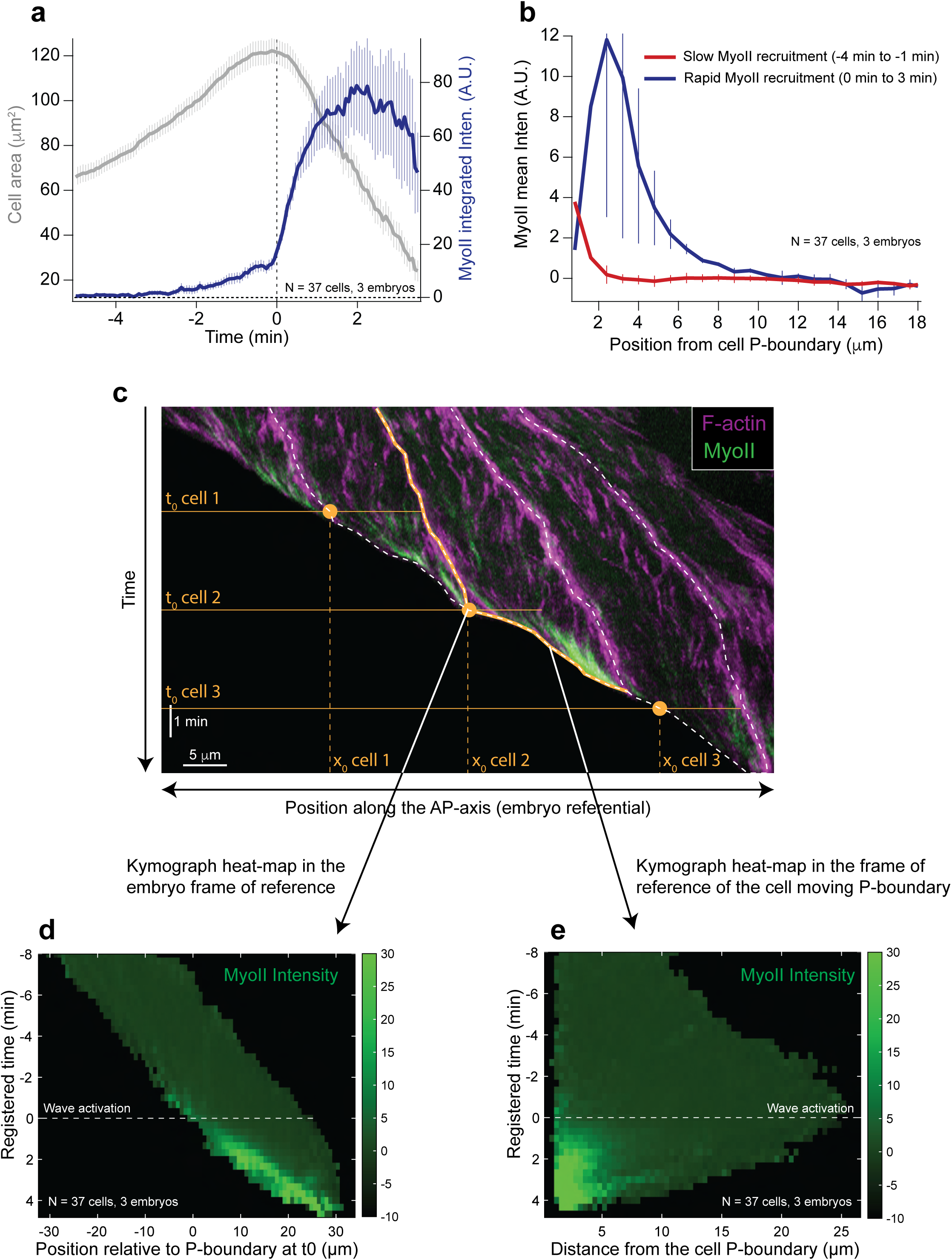
MyoII recruitment in cells and methods for cell registration. (**a**) Time trace of MyoII integrated intensity and cell area in cells of the propagation zone. Time 0 is defined for each cell as the onset of rapid MyoII recruitment (see methods). Mean ± s.e.m between cells, N=37 cells from 3 embryos. (**b**) Spatial distribution of MyoII mean intensity within the cell at the indicated time windows. Time 0 is defined as in **a**. Spatial cell registration is performed as illustrated in **c** end **e**. Mean ± SDs between embryos are shown. N=3 embryos (37 cells). (**c**) Kymograph along the AP-axis of a region of the dorsal epithelium during wave propagation. Illustrated are the 2 types of spatial cell registration used in this study. In all cases time 0 (t0) is defined as the onset of rapid MyoII recruitment (see methods). For quantifications in the fixed embryo referential (as in **d**) cells are registered spatially by defining x=0 the position of the cell posterior boundary (P-boundary) at time t=0. For quantifications in the referential of moving P-boundary of the cell (as in **e**), x=0 is the position of the cell P-boundary at each time point t. The dashed white lines highlight the A-and P-boundaries of three consecutive cells along the AP-axis. The solid orange line highlights the moving P-boundary of the middle cell. Indicated are the time t0 of the three cells. The orange solid circles indicate the position of the P-boundary at t0 for the three cells. (**d-e**) Kymograph heatmap of MyoII mean intensity in the fixed referential of the embryo (**d**) and in the moving referential of the cell P-boundary (**e**). The same data are plotted in **d** and **e**, N=37 cells, 3 embryos.

**Supplementary Figure 2:**
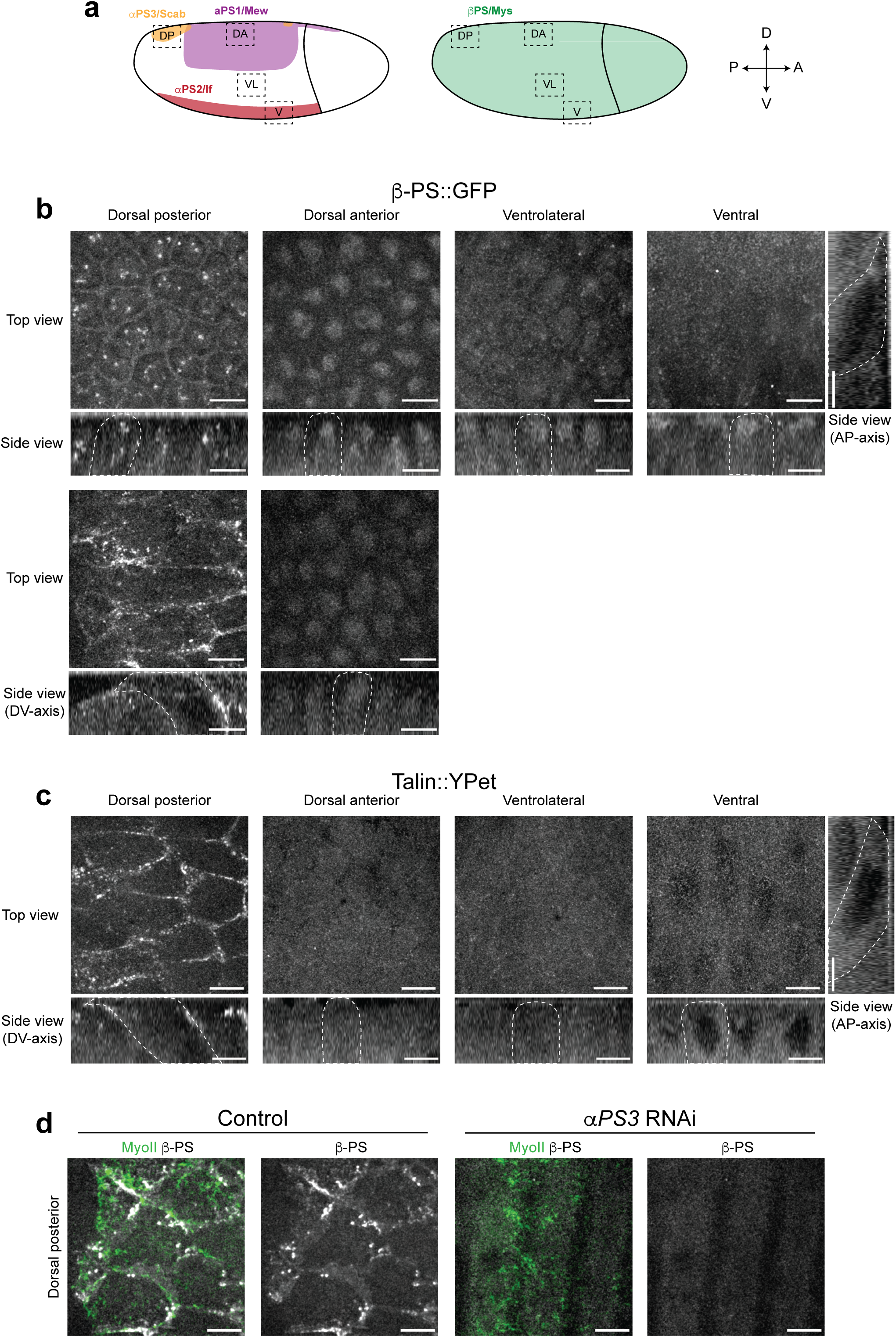
Integrin distribution in the embryo. (**a**) Cartoon representation of the expression patterns of the 5 *Drosophila* α-Integrins (left) and of β-PS (*mys*), the main β-Integrin, (right) in the embryo at gastrulation stage. The patterns were derived from Sawala et al. (Sawala et al., 2015a). The dashed boxes indicate the imaged regions in **b** and **c**. DP is dorsal posterior, DA dorsal anterior, VL ventrolateral and V is ventral. (**b-c**) Representative top and side views of β-PS (**b**) and Talin (**c**) in the indicated regions. The top views are maximum projections covering a depth of 9 µm. Side views are slices along the DV-axis and for ventral regions also slices along the AP-axis are shown. In **b** for the dorsal posterior and dorsal anterior regions images at onset of gastrulation (on the top) and mid gastrulation during wave propagation (on the bottom) are shown. All other images are at mid gastrulation stage. (**d**) Representative stills of MyoII and β-PS in the dorsal posterior region in a control (water injection) and *αPS3* RNAi injected embryo.

**Supplementary Figure 3:**
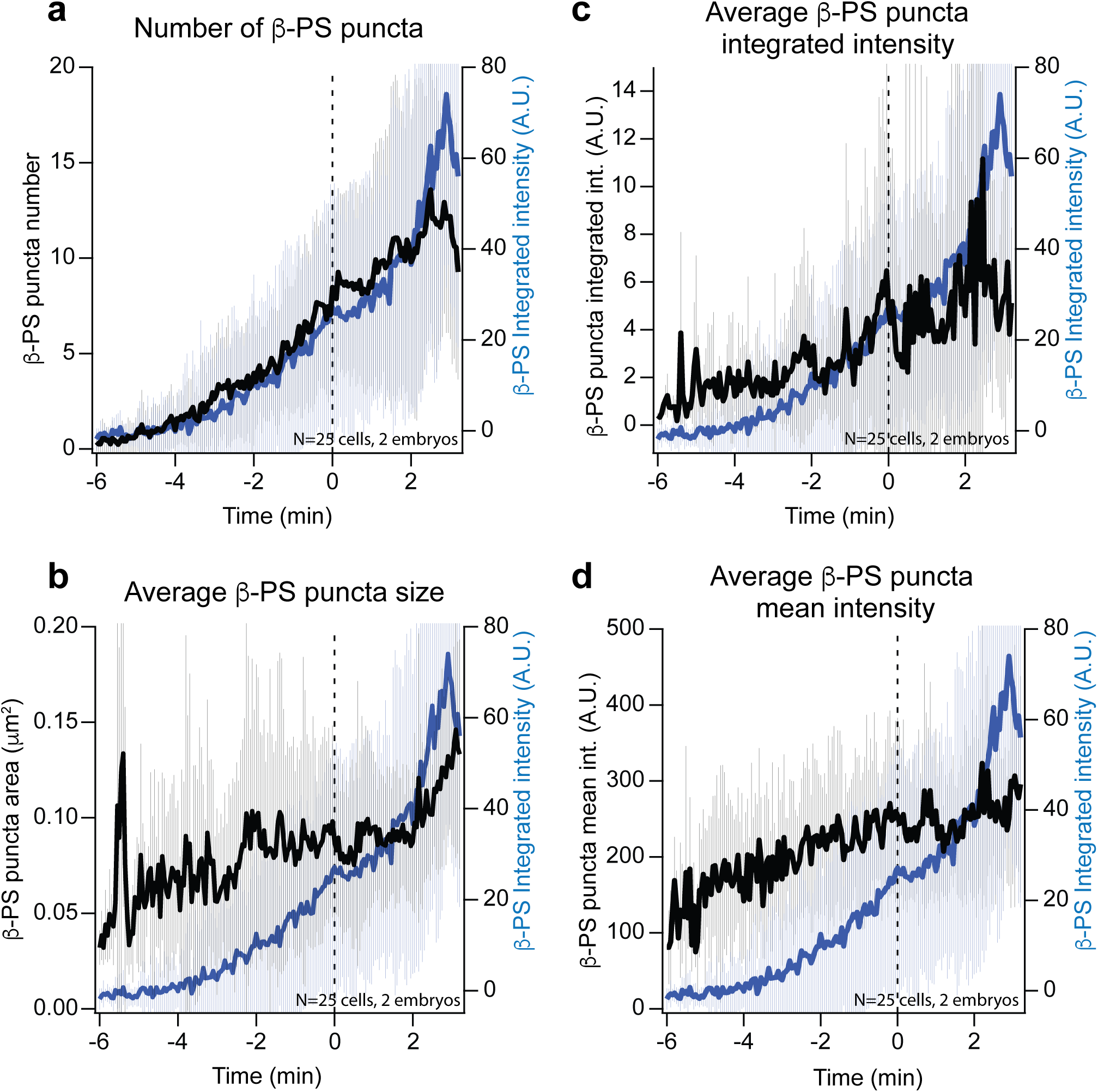
Time evolution of β-PS puncta during wave propagation. (**a-d**) In black the time traces of the average number of β-PS puncta per cell (**a**), their average size (**b**), their average integrated intensity (**c**) and their average mean intensity (**d**) together with the total β-PS integrated intensity per cell in blue. Time 0 is defined for each cell as the onset of rapid MyoII recruitment (see methods). Mean ± SDs between cells. N=25 cells from 2 embryos.

**Supplementary Figure 4:**
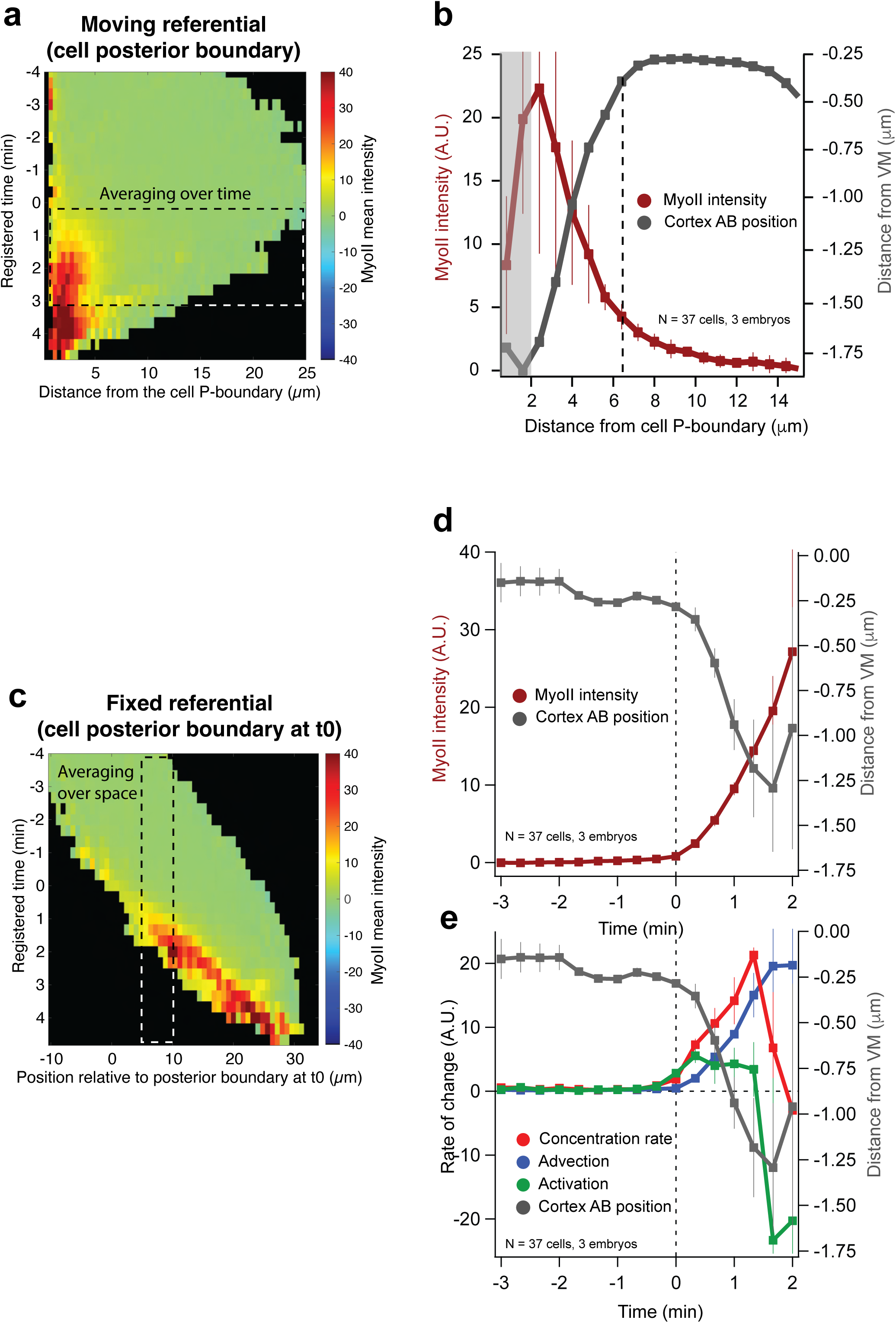
Averaging for spatial and temporal profiles of MyoII intensity and MyoII rates quantifications. (**a** and **c**) Heatmap kymograph of MyoII intensity in WT embryos. The data are plotted either relative to the moving P-boundary of cells (**a)** or in the fixed referential of the embryo (**c**). The dashed boxes indicate the region of the kymograph used to average along the time axis to obtain spatial profiles (as in **b**) or along the x-axis to obtain temporal profiles in a fixed region of the cortex (as in **d-e**). (**b**) Spatial profiles of MyoII mean intensity along with the cortex apico-basal position in WT cells during wave propagation. The averaging time window is 0-3 min as illustrated in **a**. The dashed line indicates cortex detachment and the gray box indicates where cells disappear from the field of view and measurements become less reliable. Mean ± SDs between embryos. (**d-e**) Time evolution of MyoII mean intensity (**d**) and of the MyoII rates (**e**) along with the cortex apico-basal position in a fixed region of the cortex during wave propagation in WT embryos. The averaging window is 5µm wide between position x=3µm and x=8µm, as illustrated in **b**. In all panels N=3 WT embryos (37 cells). Mean ± s.e.m between embryos.

**Supplementary Figure 5:**
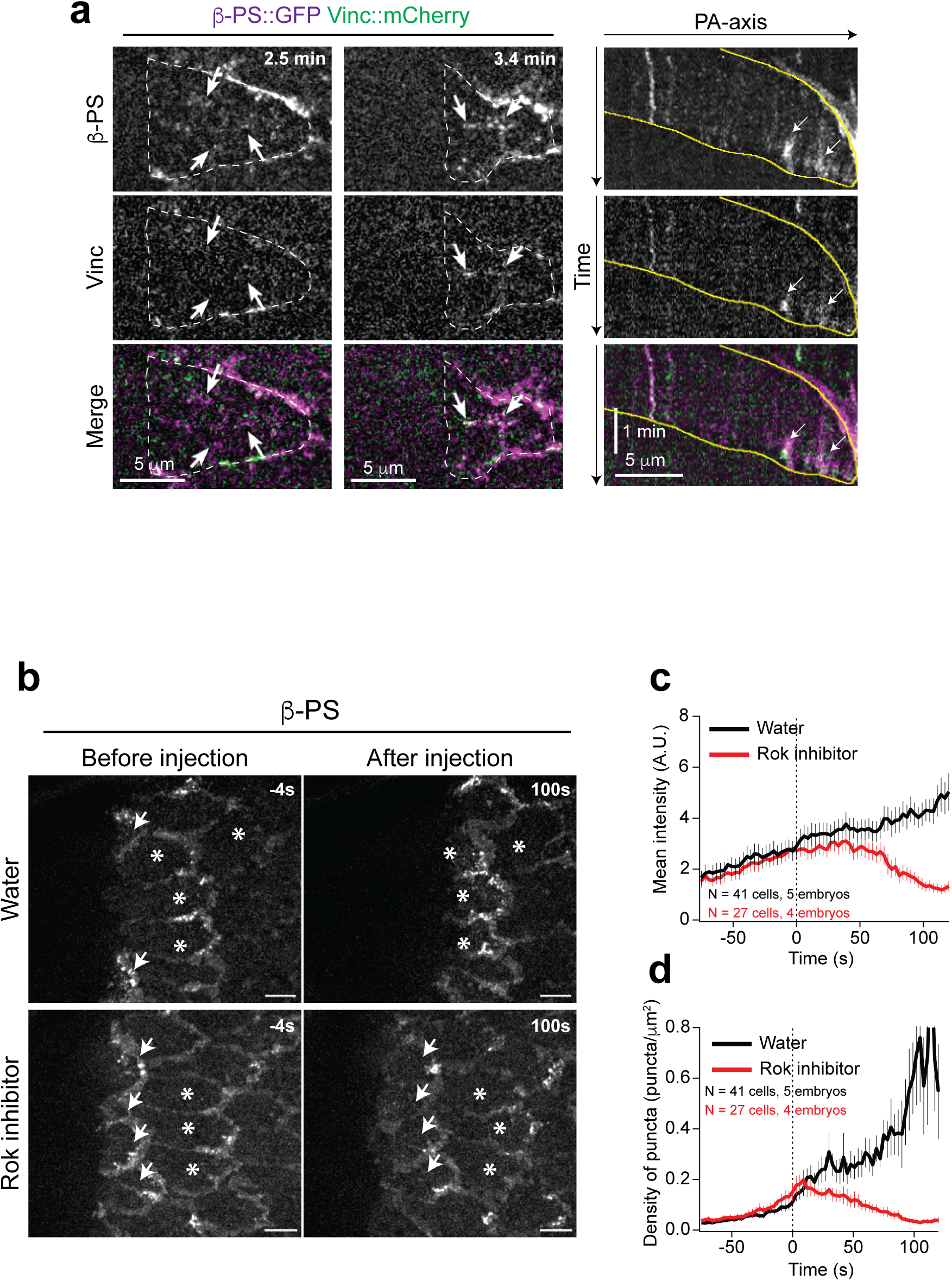
Vinculin is recruited on β-Integrin puncta and MyoII sustains their assembly. (**a**) Left: Stills from a time lapse of β-PS and Vinculin during wave propagation in a cell of the propagation zone. The dashed white line highlights the cell contour. Right: kymographs along the PA-axis of the cell on the left. The yellow lines highlight the A-and P-boundaries of the cell. The white arrows indicate β-PS puncta recruiting Vinculin. (**b**) Larger views of β-PS in the posterior endoderm before and after water or Rok inhibitor injection during wave propagation. The white arrows indicate the cells at the edge of the invaginating furrows (quantifications shown in Fig.4h-i) and the white asterisks cells where MyoII increases slowly and the intracellular wave has not yet been activated at the time of injection. (**c-d**) Quantifications of β-PS mean intensity (**c**) and β-PS density of puncta (**d**) over time in cells where MyoII increases slowly (white asterisks in **b**). Mean ± s.e.m between cells, N=41 cells from 5 water injected embryos in and N=27 cells from 4 Rok inhibitor injected embryos. Time 0 is the time of injection.

**Suppl. Fig. 6:**
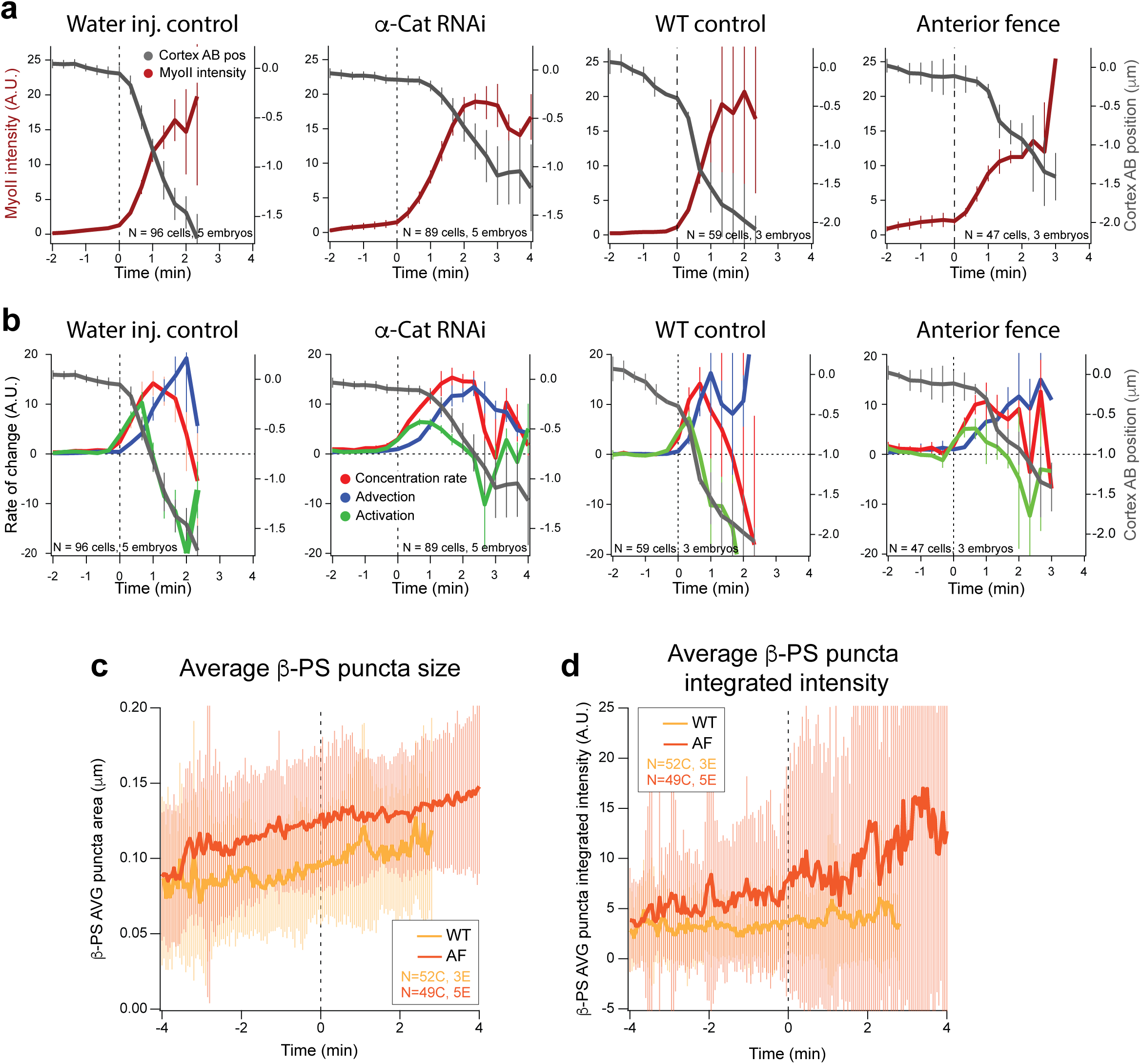
Time evolution of MyoII Intensity and MyoII rates when detachment is impaired. (**a-b**) Time evolution of MyoII mean intensity (**a**) and of the MyoII rates (**b**) along with the cortex apico-basal position in a fixed region of the cortex in the indicated conditions. N=5 embryos (96 cells) for water injection, 5 embryos (89 cells) for α-Cat RNAi, 3 embryos (59 cell) for WT control and 47 cells from 3 embryos for Anterior fence. Mean ± s.e.m between embryos. (**c-d**) Time traces of the average size (**c**) and average integrated intensity (**d**) of β-PS puncta in cells in the indicated conditions. N=52 cells from 3 WT and 49 cells from 5 embryos with anterior fences. Mean ± SDs between cells. In all cases time 0 is the time of wave activation (rapid MyoII recruitment, see methods).

**Supplementary Figure 7:**
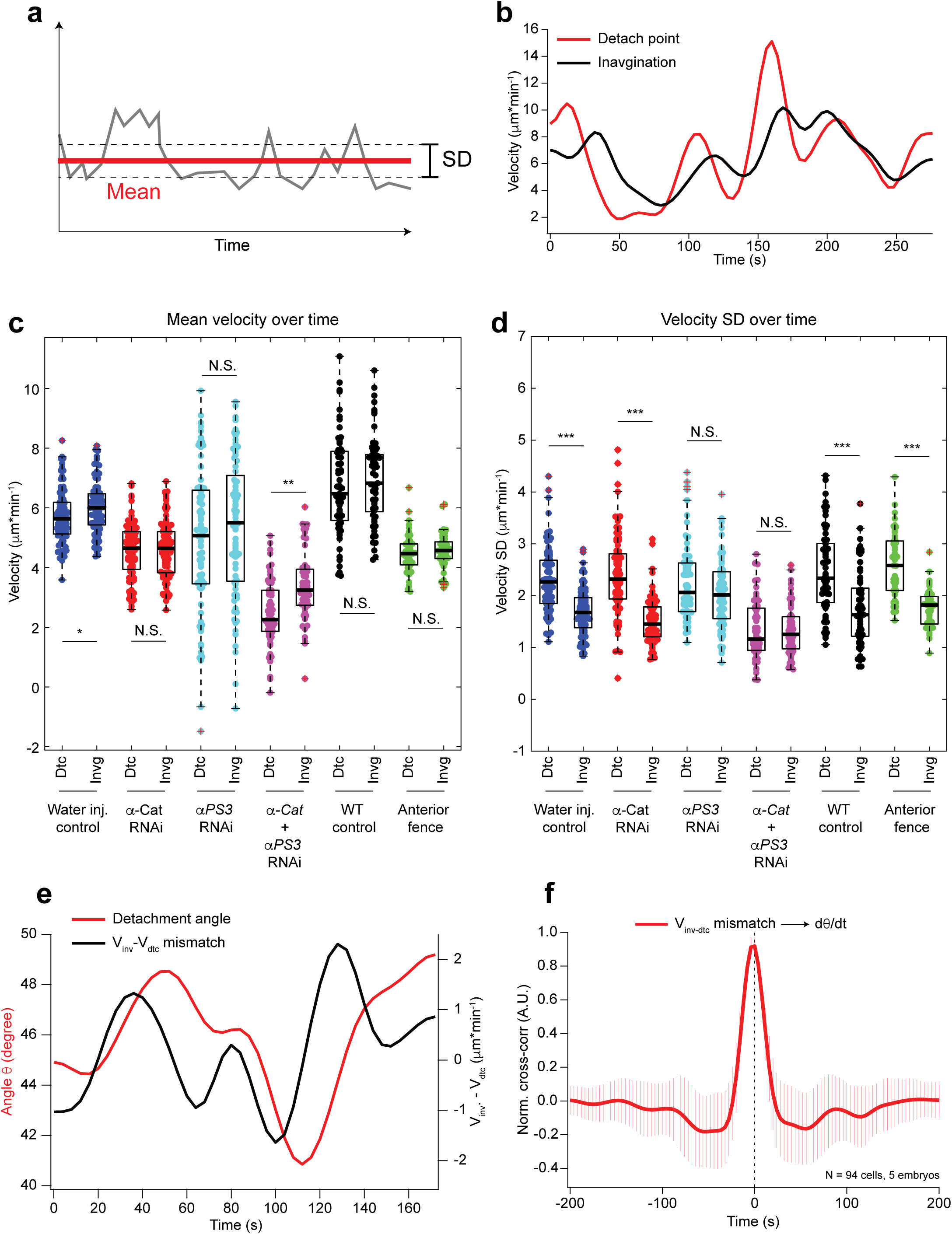
The detachment angle changes due to a mismatch between the detachment point velocity and the anterior movement of the basal furrow. (a) Illustration of a time trace (in grey) with its mean value (in red) and standard deviation (black dashed lines). The SD is used as an estimation of the variance along the time trace. (b) Representative smoothened time traces of the speed along the PA-axis of the detachment point (in red) and of the basal furrow (in black) estimated from side views as in Fig.7d. (c-d) Measurements of the mean velocity (c) and of the velocity standard deviation (d) from time traces of the speed of the detachment point (Dtc) and the basal furrow (Invg) along the PA-axis in the indicated conditions. Individual data points are superimposed to box plots. N=94 cells from 5 embryos for Water injection controls, 85 from 5 embryos for α-Cat RNAi, 76 cell from 4 embryos for *αPS3* RNAi, 59 cells from 5 embryos for *α-Cat+ αPS3* RNAi, 61 cells from 3 embryos for WT control and 47 cells from 3 embryos for Anterior fence. *** indicates a p-value <0.001, ** a p-value <0.01, * a p-value<0.05 and N.S. a p-value>0.05 at a Mann-Whitney test. (e) Representative smoothened time traces of the detachment angle θ (in red) and of the difference between the anterior velocity of the basal furrow and that of the apical detachment point (in black) in a WT embryo. (f) Average normalized cross-correlation curve between the velocity mismatch between the basal furrow the apical detachment point and the time derivatives of the detachment angle θ in WT embryos. Mean ± SDs between cells. N=94 cells from 5 WT embryos.

